# Development of human hippocampal subfield microstructure and relation to associative inference

**DOI:** 10.1101/2023.04.07.536066

**Authors:** S. Vinci-Booher, M.L. Schlichting, A.R. Preston, F. Pestilli

## Abstract

The hippocampus is a complex brain structure composed of subfields that each have distinct cellular organizations. While the volume of hippocampal subfields displays age-related changes that have been associated with inference and memory functions, the degree to which the cellular organization within each subfield is related to these functions throughout development is not well understood. We employed an explicit model testing approach to characterize the development of tissue microstructure and its relationship to performance on two inference tasks, one that required memory (memory-based inference) and one that required only perceptually available information (perception-based inference). We found that each subfield had a unique relationship with age in terms of its cellular organization. While the subiculum (SUB) displayed a linear relationship with age, the dentate gyrus (DG), *cornu ammonis* field 1 (CA1), and *cornu ammonis* subfields 2 and 3 (combined; CA2/3) displayed non-linear trajectories that interacted with sex in CA2/3. We found that the DG was related to memory-based inference performance and that the SUB was related to perception-based inference; neither relationship interacted with age. Results are consistent with the idea that cellular organization within hippocampal subfields might undergo distinct developmental trajectories that support inference and memory performance throughout development.

---

The hippocampus is a complex brain structure routinely found to be associated with memory throughout the lifespan (Scoville and Milner 1957; Scahill et al. 2003; Lister and Barnes 2009; Hirshhorn et al. 2012; Zeithamova et al. 2012; Barron et al. 2013; Schlichting et al. 2014; Venkatesh et al. 2020). The structure of the hippocampus contains distinct areas, called subfields, that are distinguishable from one another based on the properties and organization of the cells within each subfield (Manns and Eichenbaum 2006; Lavenex et al. 2007), anatomical differences that are perhaps related to subfield-specific gene expression (Datson et al. 2004; Thompson et al. 2008). Most anatomical studies relating hippocampal subfields to memory throughout the human lifespan have indexed the cellular properties of subfields by measuring their volumes (Lee et al. 2014; Pereira et al. 2014; Riggins et al. 2015; Daugherty et al. 2017; Schlichting et al. 2017). Diffusion imaging provides more specific measures of cellular properties than volume that can index changes in the orientation and density of neurons within a particular subfield (Callow et al. 2020). Yet the developmental trajectories of diffusion-derived metrics of cellular organization in hippocampal subfields and their relationships to memory throughout the lifespan are unknown. Here, we used diffusion imaging to characterize the relationship between subfield microstructure and age as well as the relationship between subfield microstructure and memory through middle childhood and young adulthood.

The volume of the hippocampus and its subfields changes throughout the lifespan (Lee et al. 2014; Pereira et al. 2014; Riggins et al. 2015; Daugherty et al. 2017; Schlichting et al. 2017). In childhood, hippocampal volume displays clear age-related increases (Hamer et al. 2008) with sex-specific trends (Giedd et al. 1996; Ostby et al. 2009). However, there is less consensus on the trajectory of hippocampal volumes in adolescence and young adulthood with some studies reporting increases, others reporting decreases, and still others reporting no change (Lee et al. 2017). Measuring age-related changes in the volume of the entire hippocampus may mask underlying trends in its anatomically distinct subfields (**Figure 1A**). Indeed, studies that investigated the development of particular hippocampal subfields, as opposed to differences across the entire hippocampus, revealed that the volumes of each subfield undergo distinct developmental trajectories from childhood through young adulthood (Lee et al. 2014, 2017; Daugherty et al. 2017; Tamnes et al. 2018). While CA1 volume displays a non-linear inverted U-shaped trajectory, CA2/3 and DG volumes display linear downward trajectories, and SUB volume displays a more complex trajectory with an early peak and a late valley. Thus, the hippocampal subfield volumes each have distinct developmental trajectories, suggesting that the nature of their cellular changes might also be distinct.

**Figure 1.**
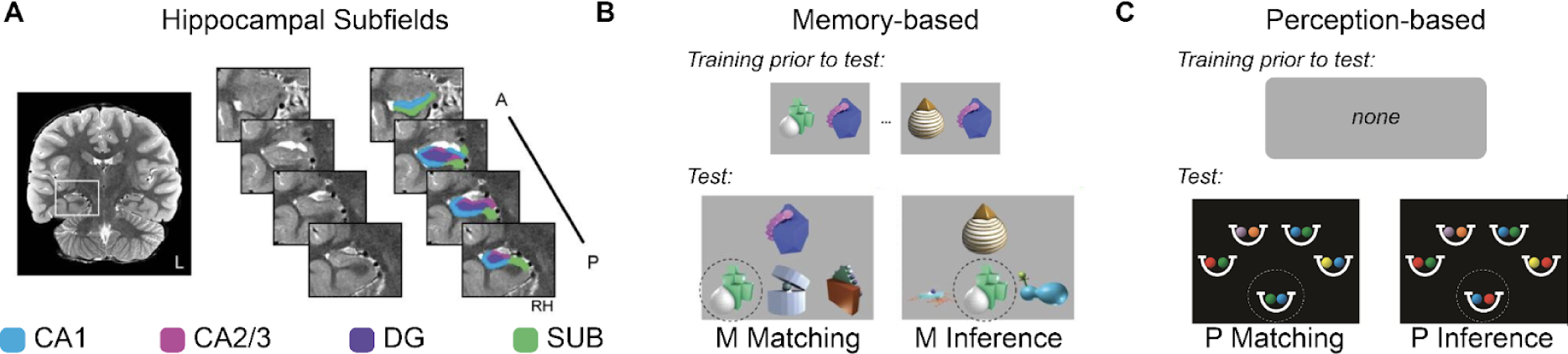
Hippocampal subfields and behavioral tasks. **A.** Hippocampal subfields. The hippocampus was segmented into four subfields: CA1 (blue), CA2/3 (pink), DG (purple), and SUB (green). **B. Memory-based inference.** In the memory-based matching and inference tasks, participants learned pairs of objects and were later required to select the object from an array of three objects that they learned with a target object (M Matching) and to select the object from an array of three objects that should go with the target object based on a shared feature (M Inference). **C. Perception-based inference.** In the perception-based matching and inference tasks, participants were asked to indicate whether or not the target set of balls (emphasized here with a circle) was a valid pair based on the other pairs of balls present on the screen (P Matching) and based on a shared feature (P Inference). Images A and B were adapted from (Schlichting et al. 2017); image C was adapted from (Wendelken and Bunge 2010). CA1 = *cornu ammonis* field 1, CA2/3 = *cornu ammonis* fields 2 and 3 combined, DG = dentate gyrus, SUB = subiculum, A = anterior; P = posterior.

An emerging body of work is using diffusion imaging to provide information about the development of cellular organization within the hippocampus and its relationship to memory. Diffusion-derived metrics of the hippocampus have developmental trajectories and behavioral correlates that are independent from their volumes (Lee et al. 2014; Pereira et al. 2014; Wolf et al. 2015; Callow et al. 2020; Langnes et al. 2020). For example, mean diffusivity (MD), an index of cellular density (Basser et al. 1994; Pierpaoli and Basser 1996; Assaf and Pasternak 2008; Le Bihan 2014), within the hippocampus decreased from 4 years to 13 years of age (Mah et al. 2017; Callow et al. 2020) and was related to performance on a source memory task (Callow et al. 2020), even after controlling for hippocampal volume. In a lifespan study of anterior and posterior hippocampal regions, results indicated different developmental trajectories for MD and volume, with region-specific trends in MD and age-related relationships between region MD and recall memory (Langnes et al. 2020). Investigations of diffusion metrics in hippocampal *subfields* in healthy, non-geriatric populations are still rare to non-existent. Consequently, the developmental trajectories of cellular organization within subfields and their relationships to memory development are nearly entirely based on volume measurements.

Evidence from mouse models suggests that measuring fractional anisotropy (FA), a diffusion-derived metric, may provide important information about the cellular organization within the hippocampus. First, histological work revealed that hippocampal subfields have characteristic fiber orientations. The predominant orientation of diffusion is dorsal-ventral in the DG, rostral-caudal in the CA3 subfield, and either dorsal-ventral or rostral-caudal depending on location within the CA1 subfield (Sierra et al. 2015). Second, experience-induced changes in fiber orientation within subfields can be captured using diffusion-derived estimates of FA. Epileptic injury resulted in an increase in the number of fibers with similar orientations (i.e., a move towards more cellular organization) in the DG subfield as well as two subareas of the CA3 subfields, which was consistent with an observed increase in FA (Sierra et al. 2015; Salo et al. 2017). Third, experience-dependent changes in cellular fiber orientations (indexed by FA) can occur within hippocampal subfields with little to no changes in neuronal density (indexed by MD), suggesting that FA and MD are indexing different tissue properties. While epileptic injury resulted in an increase in FA within DG and CA3bc, there was no observable change in MD in either subfield (Sierra et al. 2015). Furthermore, human MRI studies have demonstrated the feasibility and utility of using FA to measure cellular changes within human hippocampal subfields. FA can be measured reliably in the human hippocampus using diffusion imaging (Müller et al. 2006) and provides information about cellular organization that is independent of MD (Shereen et al. 2011; Treit et al. 2018). FA of the entire hippocampus changes with age (Carlesimo et al. 2010; Venkatesh et al. 2020) and with spatial navigation learning in adult humans (Iaria et al. 2008), suggesting that, as in mouse models, changes in the FA of hippocampal subfields may be related to behavior and experience.

Memory allows us to not only reflect upon past experiences, but also to connect related memories in order to derive new knowledge—a faculty termed memory integration (Schlichting and Preston 2015; Morton et al. 2017). Memory integration is thought to underlie successful associative inference because new memories are encoded in the context of prior memories. As such, encoded neural representations include both the directly experienced and inferential associations, and would thereby facilitate successful inference. However, memory integration may not support successful inference in childhood (Bauer and San Souci 2010; Bauer et al. 2020; Schlichting et al. 2022). Rather than integrating memories during encoding, children may encode memories separately such that successful inference in childhood would require successful memory encoding and successful reasoning about the relationships among separate memory traces. Taken together, the neural mechanisms supporting successful inference in adulthood may be qualitatively different from the neural mechanisms supporting successful inference in childhood.

Nearly all studies investigating memory integration in the hippocampus have used associative inference paradigms. Associative inference paradigms require participants to learn two independent associations among three objects, such as A goes with B and B goes with C, and they must later infer that A goes with C (Schlichting et al. 2017). Associative inference paradigms, therefore, require successful association learning of the two pairs (i.e., memory encoding) and also successful inference about the unshared features based on a shared feature (i.e., reasoning about the relationships among separate memorie encodings). Because successful inference in childhood may rely on separate processing for memory encoding and reasoning about the encoded memories, it is important to use not only standard associative inference paradigms that include memory encoding and reasoning requirements but also “perception-based” inference paradigms to assess reasoning directly (Brainerd and Reyna 1992; Crone et al. 2009). Perception-based inference paradigms require participants to make the same inference as associative inference paradigms, that A goes with C, but the inference does not require long-term memory of the two directly experienced pairs because AB and BC are both visible during inference (Wendelken and Bunge 2010). Comparing the performance on these two tasks can isolate the memory-based component of inference, given that the reasoning component should logically be similar between associative inference tasks and perception-based inference tasks.

The present work characterizes the development of the tissue microstructure of hippocampal subfields and provides information concerning the relationship of subfield tissue properties to memory and inference. We used diffusion imaging to assess the relationship between age and hippocampal subfields based on their tissue microstructure. More specifically, we employed an explicit model testing paradigm to model the relationship between the tissue microstructure of subfields CA1, CA2/3, DG, and SUB and age from 6 - 30 years. We then performed separate regressions to predict performance on two inference tasks using subfield microstructure, one that required memory (i.e., associative inference, hereafter referred to as memory-based inference; **Figure 1B**) and one did not require memory (i.e., perception-based inference, **Figure 1C**). For both regressions, we included performance on a control task. For memory-based inference, the control task was performance on memory for directly experienced pairs. For perception-based inference, the control task was essentially a matching task for associations. This approach allowed us to disentangle relationships between subfields and memory and inference throughout development. If a relationship exists between the tissue properties of a given hippocampal subfield and performance on the memory-but not perception-based inference task, it would be consistent with the notion that the cellular organization of that subfield supports memory integration processing (because memory integration is not necessary for perception-based reasoning; although see (Wimmer and Shohamy 2012; Munnelly and Dymond 2014). Such a subfield may facilitate memory-based inference by connecting directly experienced pairs in memory, rather than through a separate reasoning-based inference process.

## Materials and methods

### Participants

A total of 90 participants were recruited for this study, originally reported in (Schlichting et al. 2017): 31 children (ages 6-11 years), 25 adolescents (ages 12-17 years), and 34 adults (ages 18-30). Only participants that met inclusion several exclusion criteria were used for analysis: (1) absence of psychiatric conditions, i.e., Child Behavioral Check List in children (CBCL; (Achenbach and Edelbrock 1991)) and Symptom Checklist 90-Revised in adults (SCL-90-R, (Derogatis 1977)) in the normal range and (2) average intelligence, i.e., Weschler Abbreviated Scale of Intelligence, Second Edition (Wechsler 2018), full Scale IQ composite score above 2 standard deviations below the mean of a normative sample, resulting in the removal of 1 child, 1 adolescent, and 10 adults, and a remaining sample size of 78 participants (30 children, 24 adolescents, 24 adults). All procedures were conducted under the approval of the Institutional Review Board at the University of Texas at Austin. Parental consent was obtained for participants under 18 years of age. Participants received compensation for their participation.

Hippocampal segmentations for a total of 62 of these participants were acquired from https://osf.io/m87fd/; only participants with available hippocampal subfield segmentations were included in the brain and brain-behavior analyses, leaving 62 participants (22 children, 21 adolescents, 19 adults). Of the 62 participants, an additional 6 participants (1 child, 2 adolescents, 3 adults) were removed due to an absence of one of the required image types (T1, T2, and diffusion-weighted images) or substandard data quality, resulting in 56 participants: 21 children (M = 8.76 years, SD = 1.73 years, Range = [6, 11], 8F, 13M), 19 adolescents (M = 13.79 years, SD = 1.65 years, Range = [12, 17], 9F, 10M), and 16 adults (M = 23.50 years, SD = 3.31 years, Range = [18, 28], 9F, 7M).

#### Outlier removal for tissue microstructural analyses

Diffusion data from individual participants were removed if their framewise displacement (FD) for the diffusion scan was greater than 1 mm or if their data were determined to be of low signal-to-noise ratio (SNR). This resulted in the removal of two participants (**Supplementary Figure 1**). There were no participants that were removed based on low SNR (**Supplementary Figure 2**). We also identified and removed outliers for each region-of-interest (ROI) according to statistical criteria by applying the box plot rule to the raw residuals and robust weights (Frigge et al. 1989) for each of the models tested (see **Table 2**); these data points were removed selectively for particular ROIs. The final participant counts for each model are provided in **Table 2** and individual participant data are displayed for each model in **Figure 2**.

**Figure 2.**
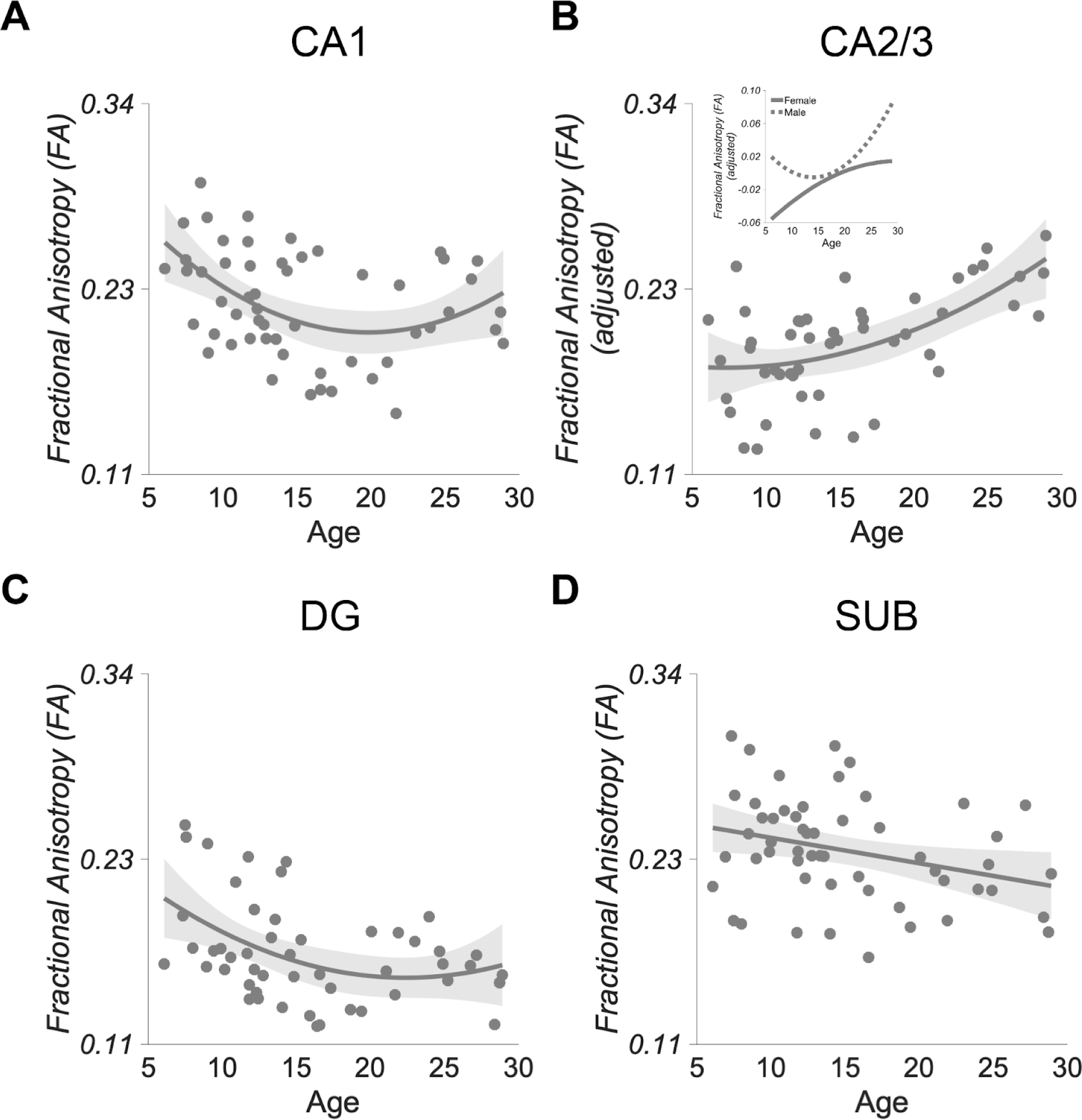
Different relationships with age for each subfield. Displayed are the best fitting models for the relationship between subfield fractional anisotropy (FA), age, and sex, selected based on *AICc*. **A.** *Cornu ammonis* field 1 (CA1). CA1 displayed a nonlinear, inverted-U shaped relationship with age. **B.** *Cornu ammonis* fields 2 and 3 (CA2/3). CA2/3 displayed a nonlinear interaction between sex and age, such that the nonlinear effect of age depended on sex. Males displayed a U-shaped relationship while females displayed the opposite. **C.** Dentate Gyrus (DG). The DG displayed a nonlinear relationship with age, such that FA decreased with age. **D.** Subiculum (SUB). The SUB displayed a linear relationship with age, such that FA decreased with age.

#### Outlier removal for behavioral analyses

Behavioral data from individual tasks performed by a participant were removed if they did not understand the task or were below chance on accuracy based on a binomial test for accuracy. The final participant counts are provided for each behavioral measure in **Figure 3** (Memory-based inference: M Matching and M Inference) and **Figure 4** (Perception-based inference: P Matching and P Inference).

**Figure 3.**
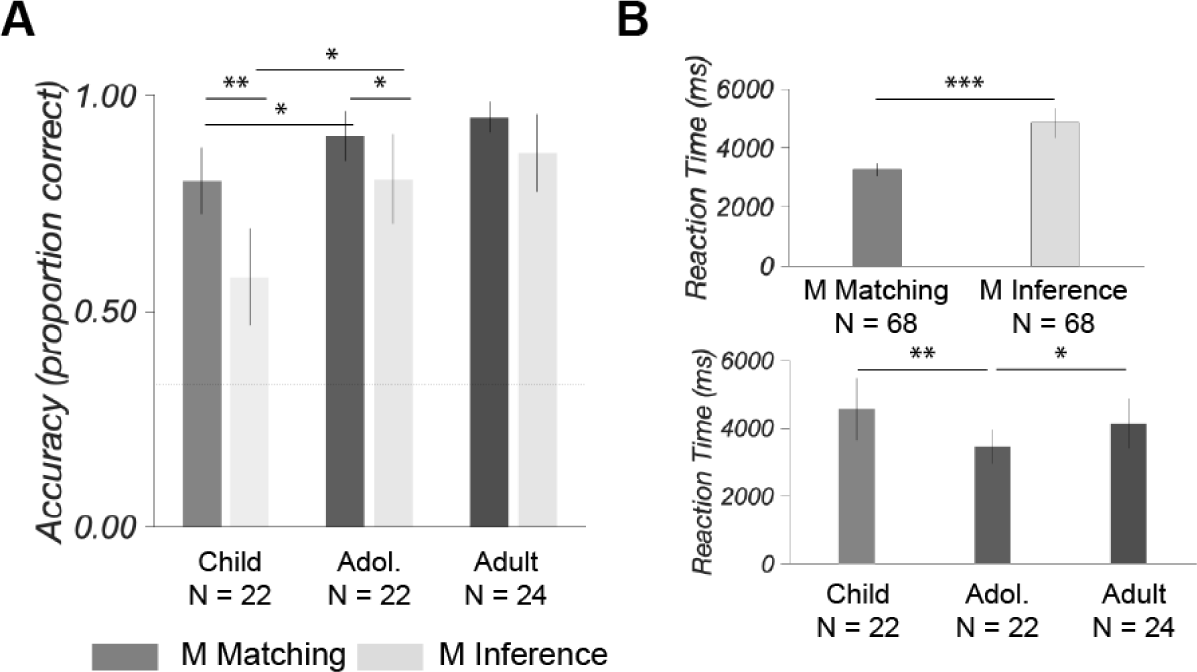
Memory-based inference: Matching and Inference. **A.** Accuracy. Performance was above chance on both tasks; chance performance was 33% (dashed line). Performance on the Matching task was greater than performance on the Inference task within the child and adolescent age groups. The difference between Matching and Inference performance was greater in children compared to adolescents. Note: This is a replication of (Schlichting et al. 2017)), Figure 4A. **B.** Reaction Time. Reaction time was slower on the Inference task than on the Matching task across age groups (top) and faster in adolescents than in children and adults across tasks (bottom). Error bars are 95% confidence intervals. *** *p* < 0.001, ** *p* < 0.01, * *p* < 0.05.

#### Outlier removal for brain-behavior analyses

We identified and removed multivariate outliers for each model. Multivariate outliers were identified using the box plot rule applied to the raw residuals as well as the robust weights (Frigge et al. 1989). The final participant counts are provided for each model along with the statistical results.

### Procedure

Participants completed two visits. During the first visit, participants completed assessments that screened for psychiatric conditions, i.e., Child Behavioral Check List in children (CBCL; (Achenbach and Edelbrock 1991)) and Symptom Checklist 90-Revised in adults (SCL-90-R; (Derogatis 1977), and potential learning disorders, i.e., Weschler Abbreviated Scale of Intelligence, Second Edition (WASI-II; (Wechsler 2018)), and were familiarized with the MRI environment in a short session in a mock MRI scanner. Participants also completed a battery of tasks designed to assess associative inference and related abilities, always completed in this order: associative inference (Preston et al. 2004; Schlichting and Preston 2015; Schlichting et al. 2017), Iowa Gambling (Bechara et al. 1994), statistical learning (Fiser and Aslin 2001; Schapiro et al. 2016; Schlichting et al. 2017), relational reasoning (Crone et al. 2009), and relational integration (Wendelken and Bunge 2010). The current work focuses on the associative inference and relational integration task because both tasks assess inference and are well matched in terms of task demands. The associative inference task assesses inference when memory is required (i.e., memory-based inference) whereas the relational integration task assesses inference when memory is not required (i.e., perception-based inference).

### Behavioral assessments

#### Memory-based inference: Matching and Inference

Participants completed an inference task that required memory. This task has been described in detail in (Schlichting et al. 2017) using stimuli that have also been described in (Schlichting and Preston 2015). A portion of the stimuli were from (Hsu et al. 2014). The task was a modified version of the task described in (Preston et al. 2004).

In brief, participants were presented with pairs of objects during a learning phase followed by a testing phase (**Figure 1B**). The 30 pairs that were split into two types of pairs: AB pairs and BC pairs. The B object was the same object for each pair type. During the learning phase, participants were encouraged to encode the object pairs by creating stories, either visual or verbal, but were told neither that the B object linked A and C objects nor that they would be asked to make an inference judgment. During the testing phase, participants were asked to complete a three-alternative forced-choice (3-AFC) task, self-paced, in which they were presented with one of the objects that they had learned and asked to identify the object that was next to it during learning, the Matching task. The cue object was presented at the top and the three choices were presented below. Distractor objects were objects that they had learned but that did not go with the cue object. Trial order was pseudo-randomized but held constant across participants and no feedback was provided. The learning and test phases were repeated 4 times each. After the 4 blocks of learning/testing, participants completed the Inference task. The Inference task was only completed once. During Inference, participants were alerted that A and C objects were linked by B items and asked to complete another self-paced 3-AFC test in which they were presented with the C object at the top and asked to select the A object from 3 objects displayed below it. All tasks were all completed using a computer and responses were provided with key presses. We will refer to the memory-based tasks as M Matching and M Inference.

#### Perception-based inference: Matching and Inference

Participants completed an inference task that did not require memory. This task was modified from the task described in (Wendelken and Bunge 2010). Note that the full set of tasks described in Wendelken and Bunge (2010) includes General Direct, General Inference, Specific Direct, and Specific Inference; however, we use only the General Direct (Matching) and General Inference (Inference) in the current work because it is so well-matched to the memory-based M Matching and M Inference tasks (described above; **Figure 1B**). In brief, participants were presented with 4 pairs of colored balls, e.g., red-blue, green-purple, yellow-white, blue-orange, and were presented simultaneously with a fifth set of colored balls, the target pair, e.g., red-orange, and asked to make a timed two-alternative forced-choice (2-AFC) judgment about the validity of the fifth set of colored balls given the associations provided in the 4 pairs (**Figure 1C**). For the Matching task, participants were able to determine the validity of the target pair by finding if that pair existed in any of the 4 pairs of colored balls. For the Inference task, participants were required to make an inference using two of the 4 pairs of colored balls presented on the screen. If, for example, a red-blue pair and a blue-orange pair exist among the 4 pairs of colored balls, then the target pair of red-orange would be considered a valid pair. Trial order was pseudo-randomized but held constant across participants and no feedback was given. We will refer to the perception-based tasks as P Matching and P Inference.

### MRI Data Acquisition and Analyses

#### MRI data acquisition

A T1-weighted 3D MPRAGE and at least two oblique coronal T2-weighted images were acquired for each participant on a 3.0T Siemens Skyra MRI. The T1-weighted image included 256 x 256 x 192 1 mm^3 voxels. The T2-weighted image was an oblique coronal acquired perpendicular to the main axis of the hippocampal complex: TR = 13150 ms, TE 82 ms, 512 x 60 x 512 matrix, 0.4 x 0.4 mm in-plane resolution, 1.5 mm thru-plane resolution, 60 slices, no gap. If visual inspection of one of the two T2-weighted images indicated motion artifacts, a third T2-weighted image was collected. Two T2-weighted images were coregistered using ANTS (Avants et al. 2011) and averaged to produce one T2-weighted image for each participant. Diffusion and T2-weighted images were aligned to the T1-weighted image.

Diffusion data were collected using single-shot spin echo simultaneous multi-slice (SMS) EPI (transverse orientation, TE = 114.00 ms, TR = 7517 ms, flip angle = 90 degrees, isotropic 1.5 mm resolution, to enable estimation of diffusion metrics within specific hippocampal subfields; FOV = LR 220 mm x 227 mm x 134 mm; Multi-band acceleration factor = 2, interleaved). Diffusion data were collected at two diffusion gradient strengths, with 64 diffusion directions at b = 1,000 s/mm^2^, as well as 1 image at b = 0 s/mm^2^, once in the AP fold-over direction (i.e., dwi-AP).

#### Anatomical (T1w) processing

Anatomical images were aligned to the AC-PC plane with an affine transformation using HCP preprocessing pipeline (Glasser et al. 2013) as implemented in the HCP AC-PC Alignment App on brainlife.io (Hayashi et al. 2018). Images from child subjects were aligned using a pediatric atlas created from 4.5-8.5 year old children (Fonov et al. 2011); images from adult subjects were aligned to the standard MNI152 adult template (Glasser et al. 2013). AC-PC aligned images were then segmented using the Freesurfer 6.0 (Fischl 2012) as implemented in the Freesurfer App on brainlife.io (Hayashi et al. 2017) to generate the cortical volume maps with labeled cortical regions according to the Destrieux 2009 atlas (Destrieux et al. 2010).

#### Diffusion (dMRI) processing

All diffusion preprocessing steps were performed using the recommended MRtrix3 preprocessing steps (Ades-Aron et al. 2018) as implemented in the MRtrix3 Preprocess App on brainlife.io (McPherson 2018). PCA denoising and Gibbs deringing procedures were performed first. We then corrected for susceptibility- and eddy current-induced distortions as well as inter-volume subject motion by interfacing with FSL tools via the *dwipreproc* script. The eddy current-corrected volumes were then corrected for bias field and Rician noise. Finally, the preprocessed dMRI data and gradients were aligned to each participant’s ACPC-aligned anatomical image using boundary-based registration (BBR) in FSL (Greve and Fischl 2009).

#### Identification of hippocampal subfields

Hippocampal subfields were identified by hand on each participant’s mean coronal image as described in (Schlichting et al. 2017, 2019). Subfields examined included: *cornu ammonis* fields 1 (CA1) and fields 2 and 3 (CA2/3), dentate gyrus (DG), and subiculum (SUB). Each subfield was segmented along the entire long axis of the hippocampus so that the microstructural measurements used in our analyses would correspond to entire subfields. Subfield segmentations included the head and body but not the tail because segmentation in the posterior part of the hippocampus was not reliable. For supplemental analyses, we further considered each subfield within head and body subregions, resulting in 8 regions of interest for the supplemental analyses: CA1_head, CA1_body, CA2/3_head, CA2/3_body, DG_head, DG_body, SUB_head, SUB_body.

#### Microstructural measurements

We estimated fractional anisotropy (FA) and mean diffusivity (MD) based on the tensor model (Basser et al. 1994) using the FSL DTIFIT App on brainlife.io (<u>brainlife.app.137</u>). Values for each microstructural measurement were extracted from hippocampal subfields using the Extract diffusion metrics inside ROIs (DTI) App on brainlife.io (<u>brainlife.app.283</u>).

### Data and software availability

Data, description of analyses, and web-links to the open source code and open cloud services used in this study are listed in **Table 1** and can be viewed in their entirety here: https://doi.org/10.25663/brainlife.pub.33. The raw data and hippocampal segmentations were transferred to the referenced brainlife.io project from the Open Science Framework (OSF): https://osf.io/m87fd/. Customized code for statistical analyses is available here: https://github.com/svincibo/devti_devHPCsubfields.

**Table 1.** Analysis steps

| Application | Github repository | Open Service DOI |
| --- | --- | --- |
| SNR Calculation | https://github.com/davhunt/app-snr_in_cc | https://doi.org/10.25663/brainlife.app.120 |
| dMRI Preprocessing | https://github.com/brain-life/app-mrtrix3-preproc | https://doi.org/10.25663/bl.app.68 |
| FSL DTIFIT | https://github.com/brainlife/app-fslDTIFIT | https://doi.org/10.25663/brainlife.app.137 |
| Extract diffusion metrics | https://github.com/brainlife/app-extract-diffusion-metrics-rois | https://doi.org/10.25663/brainlife.app.283 |

### Statistical analyses

#### Modeling the relationship between age and subfield microstructure

To determine the relationship between age and hippocampal microstructure, we assessed the relationship between hippocampal microstructure, age, and sex by testing seven different models. Models were tested for the tissue microstructure of each of the subfields (i.e., CA1, CA2/3, DG, SUB), for a total of 28 models. For each model, the dependent variable was always the mean fractional anisotropy (FA) of voxels included in the region of interest (ROI), collapsed across hemispheres. Models included: (1) ROI ∼ age, (2) ROI ∼ age^2^, (3) ROI ∼ sex, (4) ROI ∼ sex + age, (5) ROI ∼ sex + age^2^, (6) ROI ∼ sex * age, (7) ROI ∼ sex * age^2^. We included sex in our models because subfield volume has been shown to depend on sex (Tamnes et al. 2018). Continuous variables were mean centered. We identified and removed multivariate outliers for each model separately. Multivariate outliers were identified using the box plot rule applied to the raw residuals as well as the robust weights (Frigge et al. 1989). We compared the seven models for each ROI using the Akaike Information Criterion corrected for sample size (*AICc*) to account for the different numbers of predictors in each model (Hurvich and Tsai 1991). All *AICc* values were negative and we report the absolute values of *AICc* for clarity. Therefore, the model with the highest *AICc* was selected as the winning model.

Supplementary analyses included testing a series of different ROIs: hippocampal subregions (i.e., head, body, tail; instead of subfields), hippocampal subfields within head and body subregions (i.e., CA1_head, CA1_body, CA2/3_head, CA2/3_body, DG_head, DG_body, SUB_head, SUB_body), overall hippocampus. We also tested mean diffusivity (MD) rather than FA. We note that MD is commonly used in subcortical gray matter regions as well as in the hippocampus (Basser and Jones 2002).

#### Analysis of behavioral data

Each behavioral measure was analyzed separately using Two-way Repeated Measures ANOVAs with two factors: Age and Task. Age always had three levels: children, adolescents, adults. Task always had two levels. Task had two levels for the memory-based tasks, Matching and Inference; Task had two levels for the perception-based tasks, Matching and Inference. The dependent variable was either accuracy or reaction time. Pearson correlations were used to evaluate the presence of a speed-accuracy trade-off for each task (**Supplementary Figure 3**). One-sample *t*-tests were performed to confirm that performance was above chance for each task within each age group. Chance was 33% for the memory-based tasks and 50% for perception-based tasks.

#### Determining the relationships among inference, memory, and the microstructure of hippocampal subfields

To determine the relationship between the tissue microstructure of hippocampal subfields and memory performance as a function of age, we used multiple linear regression to assess the degree to which task performance on each task could predict subfield microstructure. In order to compare memory-based and non-memory based inference, we fit separate models for each subfield ROI that included a memory-based (M Matching and M Inference) and a non-memory-based task (P Matching and P Inference). Given the degree of correlation between tasks within the memory-based tasks and the non-memory-based tasks (**Supplementary Table 1**), we conducted two separate regressions for each ROI, one that included the matching tasks (M Matching and P Matching) and a second that included the inference tasks (M Inference and P Inference). We also included the model parameters found in the prior analysis that best explained the relationship between that subfield’s microstructure, age, and sex. Finally, we included IQ as a covariate of no interest to account for general intelligence, as in prior work (Schlichting et al. 2017). While accuracy and reaction time for the final repetition on the memory-based Matching task was evaluated in the behavioral analyses estimating learning, we found that the high performance on the final repetition of the memory-based Matching task left the measurement with a low level of variability and, therefore, elected to use the accuracy on the memory-based Matching task averaged across all four repetitions, as in prior work (Schlichting et al. 2017). Multivariate outliers were identified and removed for each model separately. Multivariate outliers were identified using the box plot rule applied to the raw residuals as well as the robust weights (Frigge et al. 1989). Continuous variables were mean centered. Models were considered significant with *p* < 0.025, after a Bonferroni correction for the two comparisons. Predictors within a model were considered significant with *p* < 0.05.

Statistical analyses for the repeated-measures ANOVAs that were performed on the behavioral data were performed using SPSS Statistics 27. All other statistical analyses were performed using customized code written in Matlab R2021b v.9.11.0 (see **Data and Software Availability**).

## Results

### Modeling the relationship between Fractional Anisotropy and Age: Subfields demonstrate different relationships with age

To identify developmental trends in tissue microstructure within each hippocampal subfield, we segmented the hippocampus into four subfields (i.e., CA1, CA2/3, DG, SUB) and calculated the mean fractional anisotropy (FA) as a measure of tissue microstructure for each. We then applied an explicit model testing approach to determine the best model for the relationship between FA, age, and sex. In each model, FA was entered as the response variable and age and sex were entered as predictors in one of seven possible configurations in order to test both linear and nonlinear models as well as main effect and interaction models. Pearson correlations among all variables are presented in **Supplementary Table 1**. Results of the model testing for each hippocampal subfield are displayed in **Table 2**.

**Table 2.** Models tested for Hippocampal Subfield and Age Relationship

| *Subfield* | *Regression Model* | *N* | *AICc* | *Adj. R<sup>2</sup>* | *F* | *p* |
| --- | --- | --- | --- | --- | --- | --- |
| CA1 | CA1 ~ age | 54 | 219.35 | 0.019 | 2.00 | 0.163 |
|  | **CA1 ~ age<sup>2</sup>** | **54** | **221.31** | **0.152** | **5.76** | **0.006** |
|  | CA1 ~ sex | 51 | 219.71 | 0.095 | 6.26 | 0.016 |
|  | CA1 ~ sex + age | 52 | 218.16 | 0.058 | 2.57 | 0.087 |
|  | CA1 ~ sex + age<sup>2</sup> | 55 | 221.32 | 0.107 | 3.15 | 0.033 |
|  | CA1 ~ sex * age | 54 | 217.25 | 0.081 | 1.92 | 0.139 |
|  | CA1 ~ sex * age<sup>2</sup> | 55 | 217.00 | 0.081 | 1.95 | 0.103 |
| CA2/3 | CA2/3 ~ age | 50 | 197.30 | 0.117 | 7.49 | 0.117 |
|  | CA2/3 ~ age<sup>2</sup> | 51 | 196.62 | 0.114 | 4.20 | 0.021 |
|  | **CA2/3 ~ sex** | 53 | 193.86 | 0.084 | 5.76 | 0.020 |
|  | CA2/3 ~ sex + age | 49 | 201.37 | 0.277 | 10.20 | 0.0002 |
|  | CA2/3 ~ sex + age<sup>2</sup> | 49 | 200.83 | 0.288 | 7.46 | 0.0004 |
|  | CA2/3 ~ sex * age | 49 | 198.99 | 0.260 | 6.64 | 0.0008 |
|  | **CA2/3 ~ sex \* age<sup>2</sup>** | **49** | **203.16** | **0.371** | **6.66** | **0.0001** |
| DG | DG ~ age | 51 | 209.61 | 0.114 | 7.42 | 0.009 |
|  | **DG ~ age<sup>2</sup>** | **50** | **210.00** | **0.148** | **5.27** | **0.009** |
|  | DG ~ sex | 51 | 205.87 | -0.0202 | 0.012 | 0.913 |
|  | DG ~ sex + age | 51 | 203.16 | 0.096 | 6.66 | 0.0001 |
|  | DG ~ sex + age<sup>2</sup> | 50 | 207.64 | 0.130 | 3.44 | 0.024 |
|  | DG ~ sex * age | 51 | 205.05 | 0.077 | 2.39 | 0.081 |
|  | DG ~ sex * age<sup>2</sup> | 49 | 200.75 | 0.151 | 2.71 | 0.033 |
| SUB | **SUB ~ age** | **54** | **218.07** | **0.075** | **5.30** | **0.025** |
|  | SUB ~ age<sup>2</sup> | 54 | 215.87 | 0.058 | 2.62 | 0.083 |
|  | SUB ~ sex | 55 | 216.92 | 0.011 | 1.59 | 0.213 |
|  | SUB ~ sex + age | 54 | 217.64 | 0.088 | 3.56 | 0.036 |
|  | SUB ~ sex + age<sup>2</sup> | 54 | 215.47 | 0.073 | 2.39 | 0.080 |
|  | SUB ~ sex * age | 53 | 216.31 | 0.087 | 2.66 | 0.059 |
|  | SUB ~ sex * age<sup>2</sup> | 52 | 213.41 | 0.129 | 2.51 | 0.043 |

*Best fit models are bolded, were selected on AICc, and are displayed in Figure 2.*

FA in the CA1 and DG subfields were best modeled by a nonlinear model that included only a quadratic term for age, indicating a nonlinear main effect of age for both subfields and no effect of sex. For CA1, the best model was CA1 ∼ age^2^, *AICc* = 221.31, *p* = 0.006 (**Figure 2A**). For DG, the best model was DG ∼ age^2^, *AICc* = 210.00, *p* = 0.009 (**Figure 2C**).

FA in the CA2/3 subfield was best modeled by a non-linear interaction model that included sex and age. The best model was CA2/3 ∼ sex*age^2^, *AICc* = 203.16, *p* = 0.0001, indicating the presence of a nonlinear effect of age that differed between males and females (**Figure 2B**). For males, the nonlinear effect of age was parabolic increase; for females, the effect was an inverted U-shaped relationship (**Figure 2B, inset**).

For SUB, the best model was SUB ∼ age, *AICc* = 218.07, *p* = 0.025, indicating a linear main effect of age (**Figure 2D**). As age increased, FA decreased.

Of all the additional models tested in the supplementary analyses for the hippocampus and its subregions, we found no model that fit the data better than the models tested using the subfields. In fact, nearly all models were not significant. The only exception was a simple linear model with FA in the head subregion as the response variable and sex as the predictor variable, HEAD ∼ sex, *AICc* = 255.03, *p* = 0.002. Results of these supplemental analyses can be found in **Supplementary Table 2**. Models tested with mean diffusivity (MD) as the dependent variable were similarly non-significant with the exception of TAIL ∼ age, *AICc* = 105.41, *p* = 0.005, SUB ∼ age, *AICc* = 121.91, *p* = 0.001. Results of the supplemental analyses for MD can be found in **Supplementary Table 3** for hippocampal subregions and in **Supplementary Table 4** for hippocampal subfields.

### Memory-based inference improves over development

#### Memory-based inference: Matching and Inference

To investigate memory-based inference performance as a function of age, we performed a two-way repeated-measures ANOVA for Task (Matching, Inference) and Age Group (children, adolescents, adults). The dependent variable was either accuracy or reaction time. Additionally, we used a one-sample *t*-test to ensure that accuracy was above chance, i.e., greater than 33%. We note that a prior study reported results of this analysis (Schlichting et al. 2017), and are reproduced in the current study that uses an overlapping set of participants for the benefit of the reader.

***Accuracy.*** Results revealed a greater difference between performance on the Matching task and the Inference task in children compared to adolescents (**Figure 3A**). All age groups performed above chance on both tasks, all *p*s < .001. A two-way repeated-measures ANOVA for Age Group (children, adolescents, adults) and Task (Matching, Inference) revealed a significant main effect of Age Group, *F*(2, 655) = 8.900, *p* = 3.83×10^−4^, a significant main effect of Task, *F*(1, 65) = 41.311, *p* = 1.79×10^−8^, and a significant interaction, *F*(2, 65) = 4.388, *p* = 0.016. There was a significant difference among the age groups on the Matching task, *F*(2, 67) = 6.418, *p* = 0.003, and a significant difference among age groups on the Inference task, *F*(2, 67) = 8.594, *p* = 4.88×10^−4^. For the Matching task, adult and adolescent performance were greater than child performance, *t*(44) = 3.499, *p* = 0.001 and *t*(42) = 2.077, *p* = 0.044, respectively, and there was no significant difference between adult and adolescent performance, *t*(44) = 1.288, *p* = 0216. For the Inference task, results were similar; adult and adolescent performance were greater than child performance, *t*(44) = 3.999, *p* = 2.39×10^−4^ and *t*(42) = 2.900, *p* = 0.006, respectively, and there was no significant difference between adult and adolescent performance, *t*(44) = 0.878, *p* = 0.385. Performance on the Matching task was greater than performance the Inference task in children, *t*(11) = 4.113, *p* = 0.002, and in adolescents, *t*(16) = 2.319, *p* = 0.034, but not in adults, *t*(21) = 1.962, *p* = 0.063. The difference between Matching performance and Inference performance (i.e., the difference score between Matching and Inference) was greater in children than in adolescents, *t*(42) = 2.320, *p* = 0.025, and no different in adolescents than in adults, *t*(44) = 0.365, *p* = 0.717.

***Reaction time*.** Results demonstrated that responses were slower on the Inference task than on the Matching task across age groups and that adolescents responded faster than adults and children across tasks (**Figure 3B**). A two-way repeated-measures ANOVA for Age Group (children, adolescents, adults) and Task (Matching, Inference) revealed a significant main effect of Age Group, *F*(2, 65) = 4.236, *p* = 0.019, and a main effect of Task, *F*(1, 65) = 117.057, *p* = 3.54×10^−16^. The interaction did not reach significance, *F*(2, 65) = 1.1778, *p* = 0.177. Adolescents responded faster than children, *t*(42) = 2.861, *p* = 0.007, and faster than adults, *t*(44) = 2.003, *p* = 0.051, at trend level. The difference between children and adults was not significant, *t*(44) = 1.155, *p* = 0.254. Responses were faster on the Matching task compared to the Inference task, *t*(67) = 10.779, *p* = 2.81×10^−16^.

#### Perception-based inference: Matching and Inference

To investigate perception-based inference performance as a function of age, we performed a two-way repeated-measures ANOVA for Task (Matching, Inference) and Age Group (children, adolescents, adults). The dependent variable was either accuracy or reaction time. Additionally, we used a one-sample *t*-test to ensure that accuracy was above chance, i.e., greater than 50%.

***Accuracy.*** Results demonstrated that performance was greater on the P Matching task than on the Inference task across age groups (**Figure 4A**). All age groups performed above chance on both Matching and Inference tasks, all *p*s < .001. A two-way repeated-measures ANOVA for Age Group (children, adolescents, adults) and Task (Matching, Inference) revealed a significant main effect of Task, *F*(1, 63) = 73.497, *p* = 3.57×10^−12^. Performance was greater on the Matching task than on the Inference task. Neither the main effect of Age Group, *F*(2, 63) = 2.791, *p* = 0.069, nor interaction, *F*(2, 63) = 0.918, *p* = 0.405, were significant.

**Figure 4.**
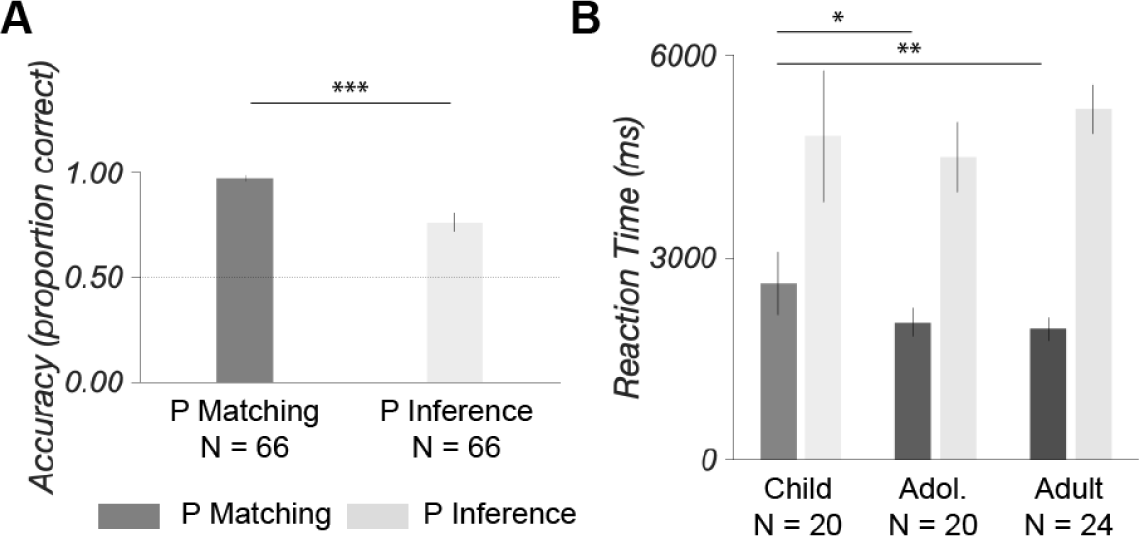
Perception-based inference: Matching and Inference. **A.** Accuracy. Performance was above chance on both tasks; chance performance was 50% (dashed line). Performance on the Inference task was less than performance on the Matching task. **B.** Reaction Time. On the Matching task, children responded slower than adolescents and adults. Error bars are 95% confidence intervals. *** *p* < 0.001, ** *p* < 0.01, * *p* < 0.05.

***Reaction time.*** Results demonstrated that children responded slower than adolescents and adults on the Matching task but not on the Inference task (**Figure 4B**). A two-way repeated-measures ANOVA for Age Group (children, adolescents, adults) and Task (Matching, Inference) revealed a significant main effect of Task, *F*(1, 63) = 316.511, *p* = 2.97×10^−26^, and a significant interaction, *F*(2, 63) = 4.746, *p* = 0.012. The main effect of Age Group was not significant, *F*(2, 63) =1.091, *p* = 0.342. Responses were slower in the Inference task than in the Matching task. There was a significant difference among age groups for the Matching task, *F*(2, 65) = 5.615, *p* = 0.006, that was not present for the Inference task, *F*(2, 65) = 1.249, *p* = 0.294. On the Matching task, children responded slower than both adolescents, *t*(42) = 2.370, *p* = 0.022, and adults, *t*(40) = 2.739, *p* = 0.009, and there was no difference between adolescents and adults, *t*(44) = 0.734, *p* = 0.467.

### Microstructure of hippocampal subfields related to associative inference

We assessed the relationship between the microstructure of hippocampal subfields and inference behavior when memory was required and when memory was not required. The goal was to explore the extent to which the cellular organization within hippocampal subfields might be related to inference behaviors by manipulating the memory requirement during inference. Associative inference requires memory and develops gradually, suggesting that the cellular computations performed within hippocampal subfields likely track memory-based inference ability and not perception-based inference ability. We constructed a separate regression model for each subfield that contained a predictor for each task of interest. One model included predictors for the inference tasks: M Inference and P Inference. The second model included predictors for the matching tasks: M Matching and P Matching. Both models included covariates of no interest, including IQ and the specific sex and age predictors that explained the most variance in subfield microstructure in the earlier analyses (see **Modelling to determine the relationship between age and subfield microstructure: Subfields demonstrate different relationships with age**). The response variable was always the mean microstructural measurement (i.e., FA) for that subfield, averaged over all voxels included in the ROI for that subfield.

#### Cornu ammonis field 1, CA1

##### Inference tasks

There was no statistically significant relationship between either of the inference tasks and CA1 microstructure. Multiple linear regression was used to test if performance on either of the inference tasks significantly predicted CA1 microstructure. The fitted regression model was CA1 ∼ M Inference + P Inference + iq + age^2^, with a final sample size of 54 after removing multivariate outliers. The overall regression was statistically significant, *R^2^* = 0.214, *Adj. R^2^* = 0.132, *AICc* = 215.98, *F*(5, 48) = 2.61, *p* = 0.036. Age significantly predicted CA1 microstructure, *β* = −0.013, *p* = 0.004, as did age^2^, *β* = 0.0004, *p* = 0.008. There were no other significant predictors, all *p*s > 0.05.

Additionally, we fitted a regression model that tested for age-related changes in the relationship between CA1 microstructure and task performance, CA1 ∼ M Inference*age + P Inference*age + iq, with a final sample size of 54 after removing multivariate outliers. However, the overall regression was not statistically significant, *R^2^* = 0.077, *Adj. R^2^* = 0.041, *AICc* = 209.44, *F*(6, 47) = 0.652, *p* = 0.688, and there were no significant predictors, all *p*s > 0.05.

##### Matching control tasks

Multiple linear regression was used to test if performance on any of the matching tasks significantly predicted CA1 microstructure. Performance on the P Matching task significantly predicted CA1 microstructure (**Figure 5A**), suggesting that more accurate performance on perception-based pair matching was related to greater directional coherence of cellular tissue within the CA1 subfield. The fitted regression model was CA1 ∼ M Matching + P Matching + iq + age^2^, with a final sample size of 53 after removing multivariate outliers. The overall regression was trending towards statistical significance, *R^2^* = 0.223, *Adj. R^2^* = 0.141, *AICc* = 217.05, *F*(5, 47) = 2.70, *p* = 0.032. P Matching significantly predicted CA1 microstructure, *β* = 0.222, *p* = 0.044, as did age, *β* = −0.012, *p* = 0.014, as did age^2^, *β* = 0.0003, *p* = 0.022. There were no other significant predictors, all *p*s > 0.05.

**Figure 5.**
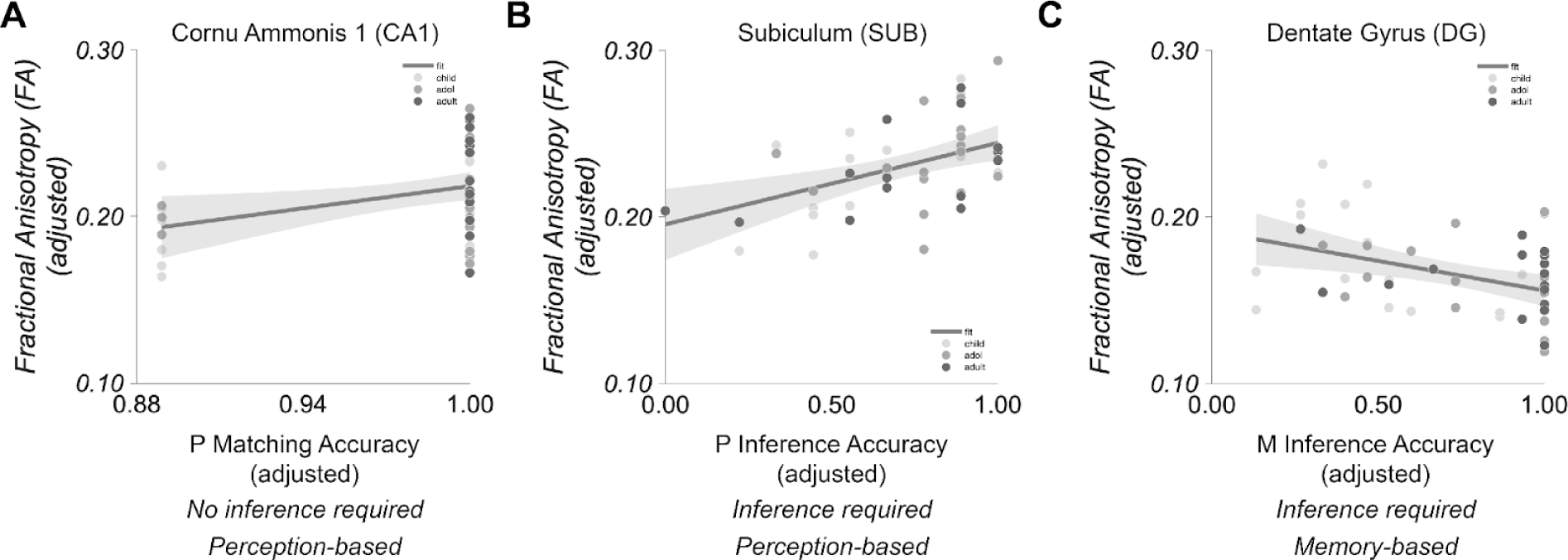
Relationship Between Subfield Microstructure and Associative Inference Performance. **A.** Perception-based matching: P Matching. P Matching accuracy significantly predicted CA1 microstructure. **B.** Perception-based inference: P Inference. P Inference accuracy significantly predicted SUB microstructure. **C.** Memory-based inference: M Inference. M Inference accuracy significantly predicted DG microstructure. Each plot describes the relationship between the fitted response as a function of accuracy with the other predictor in the model averaged out.

Additionally, we fitted a regression model that tested for age-related changes in the relationship between CA1 microstructure and task performance, CA1 ∼ M Matching*age + P Matching*age + iq, with a final sample size of 54 after removing multivariate outliers. However, the overall regression was not statistically significant, *R^2^* = 0.097, *Adj. R^2^* = −0.019, *AICc* = 210.61, *F*(6, 47) = 0.839, *p* = 0.546, and there were no significant predictors, all *p*s > 0.05.

#### Cornu ammonis fields 2 and 3, CA2/3

##### Inference tasks

There was no statistically significant relationship between either of the inference tasks and CA2/3 microstructure. Multiple linear regression was used to test if performance on any of the inference tasks significantly predicted CA2/3 microstructure. The fitted regression model was CA2/3 ∼ M Inference + P Inference + iq + sex*age^2^, with a final sample size of 48 after removing multivariate outliers. The overall regression was statistically significant, *R^2^* = 0.527, *Adj. R^2^* = 0.430, *AICc* = 199.35, *F*(8, 39) = 5.44, *p* = 0.0001. IQ significantly predicted CA2/3 microstructure, *β* = 0.0008, *p* = 0.035, as did sex, *β* = 0.177, *p* = 0.004, sex*age, *β* = −0.019, *p* = 0.016, and sex*age^2, *β* = 0.0005, *p* = 0.019. There were no other significant predictors, all *p*s > 0.05.

Additionally, we fitted a regression model that tested for age-related changes in the relationship between CA2/3 microstructure and task performance, CA2/3 ∼ M Inference*age + P Inference*age + iq, with a final sample size of 50 after removing multivariate outliers. The overall regression was statistically significant, *R^2^* = 0.245, *Adj. R^2^* = 0.140, *AICc* = 101.71, *F*(6, 43) = 2.33, *p* = 0.049. P Inference performance significantly predicted CA2/3 microstructure, *β* = −0.112, *p* = 0.041. There were no other significant predictors, all *p*s > 0.05.

##### Matching control tasks

There was no statistically significant relationship between any of the matching tasks and CA2/3 microstructure. Multiple linear regression was used to test if performance on any of the learning tasks significantly predicted CA2/3 microstructure. The fitted regression model was CA2/3 ∼ M Matching + P Matching + iq + sex*age^2^, with a final sample size of 50 after removing multivariate outliers. The overall regression was statistically significant, *R^2^* = 0.478, *Adj. R^2^* = 0.376, *AICc* = 193.44, *F*(8, 41) = 4.69, *p* = 0.0004. IQ significantly predicted CA2/3 microstructure, *β* = 0.001, *p* = 0.01, as did sex, *β* = 0.188, *p* = 0.008, and sex*age, *β* = −0.018, *p* = 0.042. There were no other significant predictors, all *p*s > 0.05.

Additionally, we fitted a regression model that tested for age-related changes in the relationship between CA2/3 microstructure and task performance, CA2/3 ∼ M Matching*age + P Matching*age + iq, with a final sample size of 53 after removing multivariate outliers. The overall regression did not reach statistical significance, *R^2^* = 0.107, *Adj. R^2^* = −0.010, *AICc* = 181.92, *F*(6, 46) = 0.916, *p* = 0.492, and there were no significant predictors, all *p*s > 0.05.

#### Dentate gyrus, DG

##### Inference tasks

Performance on the M Inference task significantly predicted DG microstructure, suggesting that the DG is related to inference that requires memory (**Figure 5C**). However, we again found no interaction with age, suggesting that the association between dentate gyrus microstructure and inference tasks may not depend on age.

Multiple linear regression was used to test if performance on any of the inference tasks significantly predicted DG microstructure. The fitted regression model was DG ∼ M Inference + P Inference + iq + age^2^, with a final sample size of 49 after removing multivariate outliers. The overall regression was statistically significant, *R^2^* = 0.420, *Adj. R^2^* = 0.353, *AICc* = 213.64, *F*(5, 43) = 6.24, *p* = 0.0002. M Inference performance significantly predicted DG microstructure, *β* = −0.036, *p* = 0.030, as did IQ, *β* = 0.0011, *p* = 0.0005. There were no other significant predictors, all *p*s > 0.05.

Additionally, we fitted a regression model that tested for age-related changes in the relationship between DG microstructure and task performance, DG ∼ M Inference*age + P Inference*age + iq, with a final sample size of 50 after removing multivariate outliers. The overall regression was statistically significant, *R^2^* = 0.406, *Adj. R^2^* = 0.323, *AICc* = 210.53, *F*(6, 43) = 4.89, *p* = 0.0007. M Inference performance significantly predicted DG microstructure, *β* = −0.082, *p* = 0.040, as did IQ, *β* = 0.001, *p* = 0.002. There were no other significant predictors, all *p*s > 0.05.

##### Matching control tasks

There was no statistically significant relationship between any of the matching tasks and DG microstructure, suggesting that the association between dentate microstructure and matching may not vary much with age.

Multiple linear regression was used to test if performance on any of the matching tasks significantly predicted DG microstructure. The fitted regression model was DG ∼ M Matching + P Matching + iq + age^2^, with a final sample size of 50 after removing multivariate outliers. The overall regression was statistically significant, *R^2^*= 0.251, *Adj. R^2^* = 0.166, *AICc* = 206.89, *F*(5, 44) = 2.95, *p* = 0.022; however, there were no significant predictors, all *p*s > 0.05.

Additionally, we fitted a regression model that tested for age-related changes in the relationship between DG microstructure and task performance, DG ∼ M Matching*age + P Matching*age + iq, with a final sample size of 50 after removing multivariate outliers. The overall regression did not reach statistical significance, *R^2^* = 0.244, *Adj. R^2^* = 0.139, *AICc* = 203.76, *F*(6, 43) = 2.32, *p* = 0.051, and there were no significant predictors, all *p*s > 0.05.

#### Subiculum, SUB

##### Inference tasks

Performance on the P Inference task significantly predicted SUB microstructure, suggesting that the SUB is related to inference that does not require memory (**Figure 5B**). Multiple linear regression was used to test if performance on any of the inference tasks significantly predicted SUB microstructure. The fitted regression model was SUB ∼ M Inference + P Inference + iq + age, with a final sample size of 49 after removing multivariate outliers. The overall regression was statistically significant, *R^2^* = 0.322, *Adj. R^2^* = 0.260, *AICc* = 216.70, *F*(4, 44) = 5.21, *p* = 0.002. P Inference performance significantly predicted SUB microstructure, *β* = 0.050, *p* = 0.003, as did age, *β* = −0.002, *p* = 0.014. There were no other significant predictors, all *p*s > 0.05.

Additionally, we fitted a regression model that tested for age-related changes in the relationship between SUB microstructure and task performance, SUB ∼ M Inference*age + P Inference*age + iq, with a final sample size of 50 after removing multivariate outliers. The overall regression did not reach statistical significance, *R^2^* = 0.228, *Adj. R^2^* = 0.120, *AICc* = 212.50, *F*(6, 43) = 2.12, *p* = 0.071, and there were no significant predictors, all *p*s > 0.05.

##### Matching control tasks

There was no statistically significant relationship between any of the matching tasks and SUB microstructure. Multiple linear regression was used to test if performance on any of the matching tasks significantly predicted SUB microstructure. The fitted regression model was SUB ∼ M Matching + P Matching + iq + age, with a final sample size of 50 after removing multivariate outliers. The overall regression was significant, *R^2^* = 0.277, *Adj. R^2^* = 0.213, *AICc* = 213.07, *F*(4, 45) = 4.31, *p* = 0.005, and one predictor reached statistical significance, M Matching, *β* = −0.071, *p* = 0.016. There were no other significant predictors, all *p*s > 0.05.

Additionally, we fitted a regression model that tested for age-related changes in the relationship between SUB microstructure and task performance, SUB ∼ M Matching*age + P Matching*age + iq, with a final sample size of 51 after removing multivariate outliers.. The overall regression was statistically significant, *R^2^* = 0.252, *Adj. R^2^* = 0.151, *AICc* = 209.44, *F*(6, 44) = 2.48, *p* = 0.038, yet no predictor reached statistical significance.

## Discussion

Our goal was to determine age-related differences in the tissue properties of hippocampal subfields and their relationships to the development of associative inference and memory. We found distinct relationships between age and cellular organization by hippocampal subfield: linear for SUB and non-linear for CA1, CA2/3, DG, that moreover depended on sex for CA2/3. We found that the tissue properties of the DG and SUB were both related to associative inference performance, controlling for age, with the DG being related to associative inference that required memory and the SUB being related to associative inference that did not require memory. Overall, our results suggest that hippocampal subfields undergo distinct developmental trajectories in terms of cellular organization which in turn have distinct relationships with associative inference and memory.

### Microstructures of hippocampal subfields demonstrate different relationships with age

We observed age-related changes in hippocampal subfields, finding that different subfields have unique relationships with age from childhood to adulthood. While CA1 and DG subfields displayed a non-linear U-shaped relationship with age, with the lowest FA in adolescents for CA1 and in adulthood for DG, the SUB displayed a linear relationship with age, with FA decreasing from childhood to adulthood. The CA2/3 subfield displayed a more complicated relationship with age, with males exhibiting a U-shaped relationship with the lowest FA in adolescents, and females exhibiting an inverted U-shaped relationship with the lowest FA in childhood.

The development of cellular organization within the hippocampus has been investigated using measures of volume across the entire hippocampus, hippocampal subregions, and hippocampal subfields (Giedd et al. 1996; Hamer et al. 2008; Ostby et al. 2009; Lee et al. 2014; Riggins et al. 2015; Daugherty et al. 2017; Schlichting et al. 2017; Tamnes et al. 2018) while investigations using diffusion imaging have focused on either the entire hippocampus or hippocampal subregions but not subfields using a single diffusion metric (i.e., mean diffusivity) (Wolf et al. 2015; Lee et al. 2017; Callow et al. 2020). In the present work we employed an additional metric, fractional anisotropy (FA), to assess cellular organization within the hippocampus. We selected this metric based on results from animal models that suggested that subfields have distinct cellular organizations that could be captured with FA (Sierra et al. 2015; Salo et al. 2017) and that FA and MD provide independent information in human hippocampus (Shereen et al. 2011; Treit et al. 2018). Our results suggest that distinct developmental trajectories for hippocampal subfields can be measured using FA. Notably, we did not find age-related changes in FA within hippocampal subregions (see **Supplementary materials**), suggesting that the changes in FA within the hippocampus might occur cohesively within subfields such that measurements of FA in subregions encompassing more than one subfield cannot adequately capture the development of FA in the hippocampus.

At least one prior study has reported age-related changes in microstructure of hippocampal subregions (Langnes et al. 2020). Using a cross-sectional design across the ages of 4 to 93 years, this study reported different age-related changes for anterior and posterior hippocampus: MD in anterior hippocampus decreases from childhood to approximately 35 years of age while MD in posterior hippocampus remains relatively stable. After the age of 35 years, MD in both anterior and posterior hippocampus increases dramatically. In the current study, we segmented the hippocampus into the head, body, and tail but found no evidence of age-related changes in these subregions overall (see **Supplemental Materials**). However, results from the FA analysis within each subfield are consistent with different age-related changes for anterior and posterior hippocampus, but suggest that age-related changes detected in the larger anterior and posterior subregions may be driven by subfield-selective changes in cellular organization, such as changes to the characteristic fiber orientations of each subfield. More research will be necessary to better understand the relationship between developmental trends in cellular organization detected in larger subregion segmentations and trends detected in smaller subfield segmentations.

### Associative inference performance that requires memory improves over development

Associative inference improves gradually throughout development (Shing et al. 2019; Schlichting et al. 2022), as do memory processes (Andrews & Halford, 1998; Bauer, Varga, King, Nolen, & White, 2015; Halford, 1984; Townsend, Richmond, Vogel-Farley, Thomas, 2010; Ghetti & Bunge, 2012). Our results further support the findings that associative inference and memory improve throughout development. Performance on an associative inference task that required memory was greater in adults compared to adolescent children and greater in adolescent children compared to younger children (originally reported in Schlichting et al., 2017), suggesting that memory-based associative inference changes throughout middle childhood and adolescence. However, we did not find a similar result for similarly structured inference task without a memory requirement: Perception-based inference performance was no different among children, adolescents, and adults. One interpretation of this null result is that associative inference in the absence of a memory requirement may reach adult-like performance earlier than 6 years of age. Although this interpretation would be consistent with reports that children are capable of perception-based reasoning with novel items by the age of 4 years old (Brainerd and Reyna 1992), it is possible that the null result was due to experimental factors. For example, the perception-based task might have been less sensitive to developmental changes than the memory-based task given that the perception-based task (i.e., 50%) had a higher floor than the memory-based task (i.e., 33%).

### Microstructures of hippocampal subfields related to associative inference and memory

Understanding the relationship between subfield development, associative inference, and memory in terms of tissue microstructure can provide insight into the nature of memory and the neural mechanisms supporting memory. We found that DG microstructure was related to associative inference performance when memory was required, but that SUB microstructure was related to associative inference performance when memory was not required. Furthermore, CA1 microstructure was related to pair matching that did not require memory. Notably, the relationships among subfield microstructure and behavior were observed after controlling for age, suggesting that age may not be a driving factor in the relationship between subfield cellular organization and associative inference and memory.

One theory of the neural mechanisms underlying associative inference proposes that the hippocampus proactively integrates new memories with existing memories through memory integration, a process of combining overlapping memory traces during encoding (Schlichting and Preston 2015). Each of the hippocampal subfields are thought to support unique aspects of the memory integration process. The DG subfield may assist memory integration by supporting memory for directly experienced pair associations through a process of pattern separation (Berron et al. 2016; Schapiro et al. 2017; Duncan and Schlichting 2018; Canada et al. 2019). While the CA3 subfield is thought to activate memories related to a novel event (McKenzie et al. 2014), the CA1 subfield is thought to use novelty as a cue to trigger the memory integration process (Larkin et al. 2014). The function of the subiculum (SUB) is less clear, however it is more densely connected to regions outside of the hippocampus than subfields in the hippocampus proper (i.e., DG, CA2/3, CA1), suggesting that it has functions that are potentially independent of core hippocampal functions (Aggleton and Christiansen 2015).

Our results suggest that hippocampal subfields may play important, yet distinct, roles in integrating new memories with existing memories during development. We observed different behavioral correlates of subfield cellular organization, namely that DG was related to associative inference that required memory while CA1 was related to pair matching that did not require memory. Although more work is needed to clarify the implications of our results, we suggest that our results support the proposal that hippocampal subfields have unique functional properties that, together, may support inference in the age range corresponding to our sample, namely middle childhood, adolescence, and young adulthood. One possibility is that DG may be important for successful inference to supply highly accurate representations of learned associations. In other words, DG microstructure might support associative inference through its ability to support accurate encoding for separate representations that would allow for more accurate reasoning about the relationships among those separate representations, a mechanism that may be more prevalent in childhood than in adulthood (Schlichting et al. 2017, 2022; Duncan and Schlichting 2018; Shing et al. 2019). While adults may support associative inference behavior by drawing on integrated hippocampal representations, children may be relying on separate, non-integrated memories for directly experienced associations. Given our developmental sample, it is possible that participants were performing the memory-based associative inference task using separate memory encodings rather than integrated memories.

Our results are consistent with the notion that the SUB may have functions that are independent of core hippocampal functions (Aggleton and Christiansen 2015). Results suggest that the cellular organization within the SUB was related to associative inference that could be accomplished based on perceptually available information and, therefore, may not necessarily be related to memory. It is possible that the SUB interfaces with other cortical mechanisms to support reasoning in the absence of relevant memories, given that it is more densely connected to regions outside of the hippocampus than the other subfields. More research will be necessary to better understand the role of the SUB in associative inference and memory.

## Conclusion

The cellular organization of hippocampal subfields demonstrate different relationships with age and exhibit distinct relations with associative inference and memory.

## Acknowledgements

The authors would like to thank Jessica Church-Lang, Michael Mack, Tammy Tran, Amelia Wattenberger, and Katie Guarino for assistance with the project, such as participant recruitment, data collection, and helpful discussions. This work was supported by the National Science Foundation (SBE Postdoctoral Fellowship SMA-2004877 to SVB; OAC-1916518, IIS-1912270, IIS-1636893, BCS-1734853 to FP; CAREER Award 1056019 to ARP), the National Institutes of Health (R01MH126699, R01EB030896, R01EB029272 to FP; R01MH100121, R21HD083785 to ARP), the Department of Defense (NDSEG Graduate Fellowship Program to MLS), the Wellcome Trust (226486/Z/22/Z to FP), and a University of Texas Research Grant to ARP. The work was also supported by a Microsoft Investigator Fellowship to FP and a gift from the Kavli Foundation to FP.

