## Supplementary materials for "Development of human hippocampal subfield microstructure and relation to associative inference"

### Supplementary analyses

#### *Supplementary analysis: Identify outliers based on motion.*

We measured the frame-wise displacement (FD) to evaluate the quality of diffusion data collected for each subject (Power et al., 2014). Two subjects were removed due to high motion, defined as FD > 1 mm (**Supplementary Figure 1A**). A One-way ANOVA indicated that there was no significant difference in the FD of diffusion images among the groups, after removing the two high motion participants, $F(2, 53) = 1.760$, $p = 0.182$ (**Supplementary Figure 1B**). A simple linear regression with FD as the response variable and Age as the predictor variable indicated no significant linear relationship between FD and Age, after removing the two high motion participants, $\beta = -7.4x10^{-4}$, $p = 0.79$ (**Supplementary Figure 1C**).

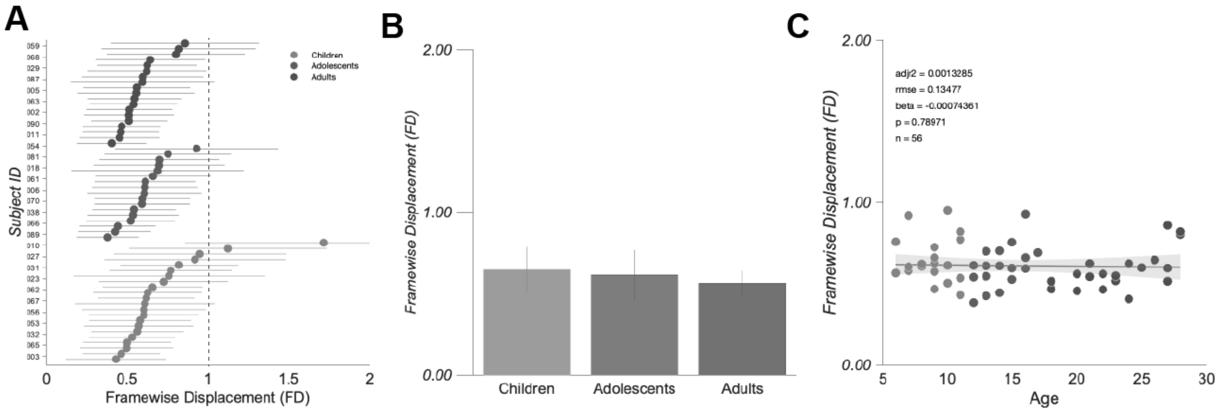


**Supplementary Figure 1.** Framewise Displacement (FD). **A.** FD for each participant, including participants that were eventually removed from certain analyses for motion or as statistical outliers. Two participants from the child group were identified as high-motion participants with FD > 1 mm (dashed line) and removed from all diffusion analyses. **B.** No significant difference in FD among age groups after removing the two high-motion participants. Mean and standard deviations are displayed for each group. **C.** No significant correlation between FD and age after removing the two high-motion participants. The line indicates the linear fit and the shaded area indicates the 95% confidence bounds.

***Supplementary analysis: Identify outliers based on SNR.***

We measured the signal-to-noise (SNR) to evaluate the quality of diffusion data collected for each participant. All SNR of all diffusion data were above 4 and no participants were removed due to low SNR (**Supplementary Figure 2A**).

A One-way ANOVA indicated that there was a significant difference in the SNR of weighted diffusion images, $F(2, 53) = 6.350$, $p = 0.003$ (**Supplementary Figure 2B**). The SNR of the child diffusion datasets was greater than both the SNR of the adult, $t(35) = 3.256$, $p = 0.002$, and than the adolescent, $t(38) = 2.161$, $p = 0.037$, datasets. The difference between adolescent and adult datasets did not reach significance, $t(33) = 1.4867$, $p = 0.147$.

A simple linear regression with SNR of the weighted volumes as the response variable and Age as the predictor variable indicated a significant linear relationship between SNR and Age, after removing the two high motion participants, $\beta = -0.309$, $p = 0.003$ (**Supplementary Figure 2C**).

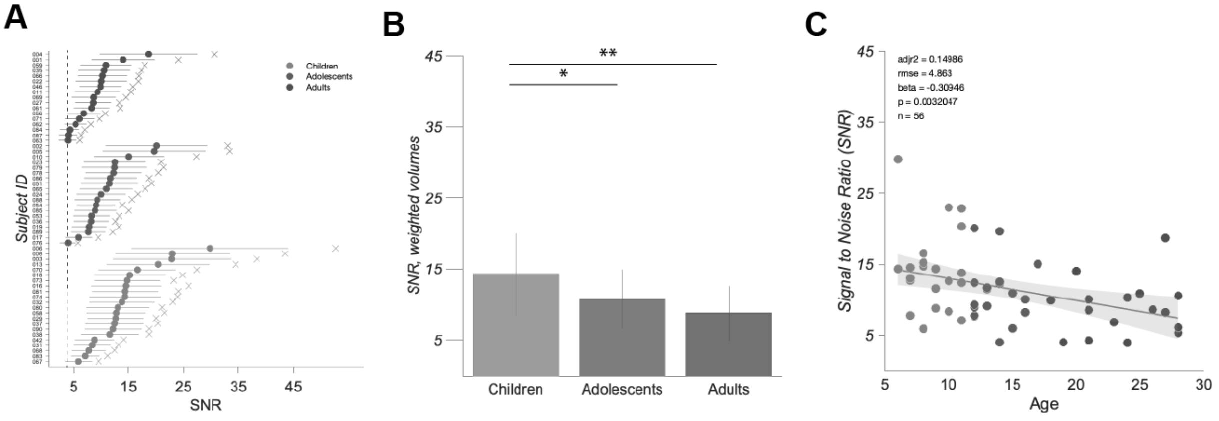


**Supplementary Figure 2.** Signal-to-noise ratio (SNR). **A.** SNR for each participant, including participants that were eventually removed from certain analyses for motion or as statistical outliers. Closed circles indicate the mean SNR across weighted diffusion images and the x marks the mean SNR of the non-weighted diffusion images, i.e., the b0 images. Lines around the mean SNR for each participant's weighted diffusion images indicate the standard deviation of the mean. SNR for all participants was greater than 4 (dashed line); no participants were removed based on low SNR. **B.** SNR was greater in children than in adolescents and adults in the diffusion weighted images. Means and standard deviations are displayed for each group. **C.** SNR of the weighted diffusion images was related to age. The linear fit and 95% confidence bounds are displayed. **, $p < 0.01$, *, $p < 0.05$.

***Supplementary analysis: Evaluation of speed-accuracy trade-offs.***

To evaluate the possibility of speed-accuracy trade-offs, we performed Pearson correlations to determine if accuracy was linearly related to reaction time in either task.

*Inference with memory requirement: ABBC Direct and AC Inference.* We observed no evidence of a speed-accuracy trade-off for the AC Inference task with a possible trade-off occurring in the ABBC Direct task (**Supplementary Figure 3A**). The correlation between accuracy and reaction time for the AC Inference task was not significant, $r = -0.032$, $p = 0.794$; however, the correlation between accuracy and reaction time for the ABBC Direct task was significant, $r = -0.419$, $p = 4\times10^{-4}$.

*Inference without memory requirement: Gen Direct and Gen Inference.* We observed no evidence of a speed-accuracy trade-off for either the Gen Inference or the Gen Direct task (**Supplementary Figure 3B**). The correlation between accuracy and reaction time for the Gen Inference task was not significant, $r = 0.183$, $p = 0.142$, and neither was the correlation between accuracy and reaction time for the Gen Direct task, $r = -0.098$, $p = 0.309$.

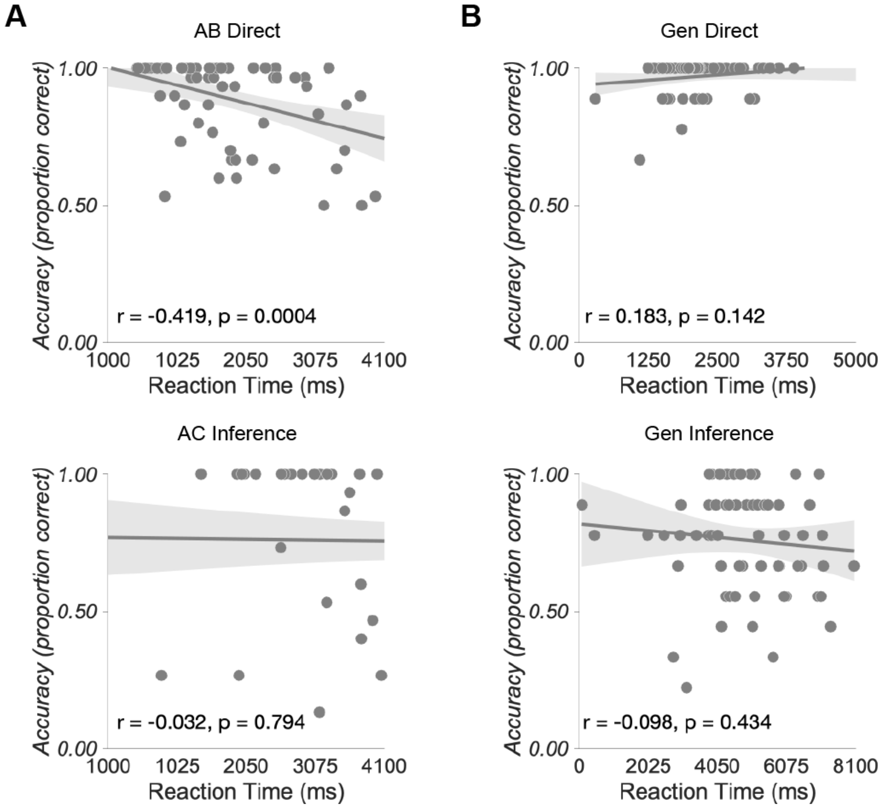


**Supplementary Figure 3.** Relationships between accuracy and reaction time for each behavioral measure. **A.** Associative inference with memory requirement: ABBC Direct (top) and AC Inference (bottom). **B.** Associative inference without memory requirement: Gen Direct (top) and Gen Inference (bottom).

***Supplementary analysis: Evaluation of diffusion measurements.***

Our FA and MD values were generally within the expected range for the hippocampus (**Supplementary Figure 4**). To evaluate the diffusion measurements, we plotted fractional anisotropy (FA) and mean diffusivity (MD) for each subfield and subregion and compared these values to the range of values reported for the hippocampus in prior works. We identified two prior works that reported on both FA and MD measurements within the hippocampus, specifically (Müller et al. 2006; Anblagan et al. 2018). In Müller et al. (2006), the FA range was approximately [0.20 0.30] and the MD range was approximately [0.70 0.80]. In Anblagan et al. (2018), the FA range was approximately [0.08 0.15] and the MD range was approximately [0.70 1.20]. We selected the lowest and highest values between the two prior works for each measurement to create the expected range of values for our dataset, resulting in an expected FA range of [0.08 0.30] and expected MD range of [0.70 1.20].

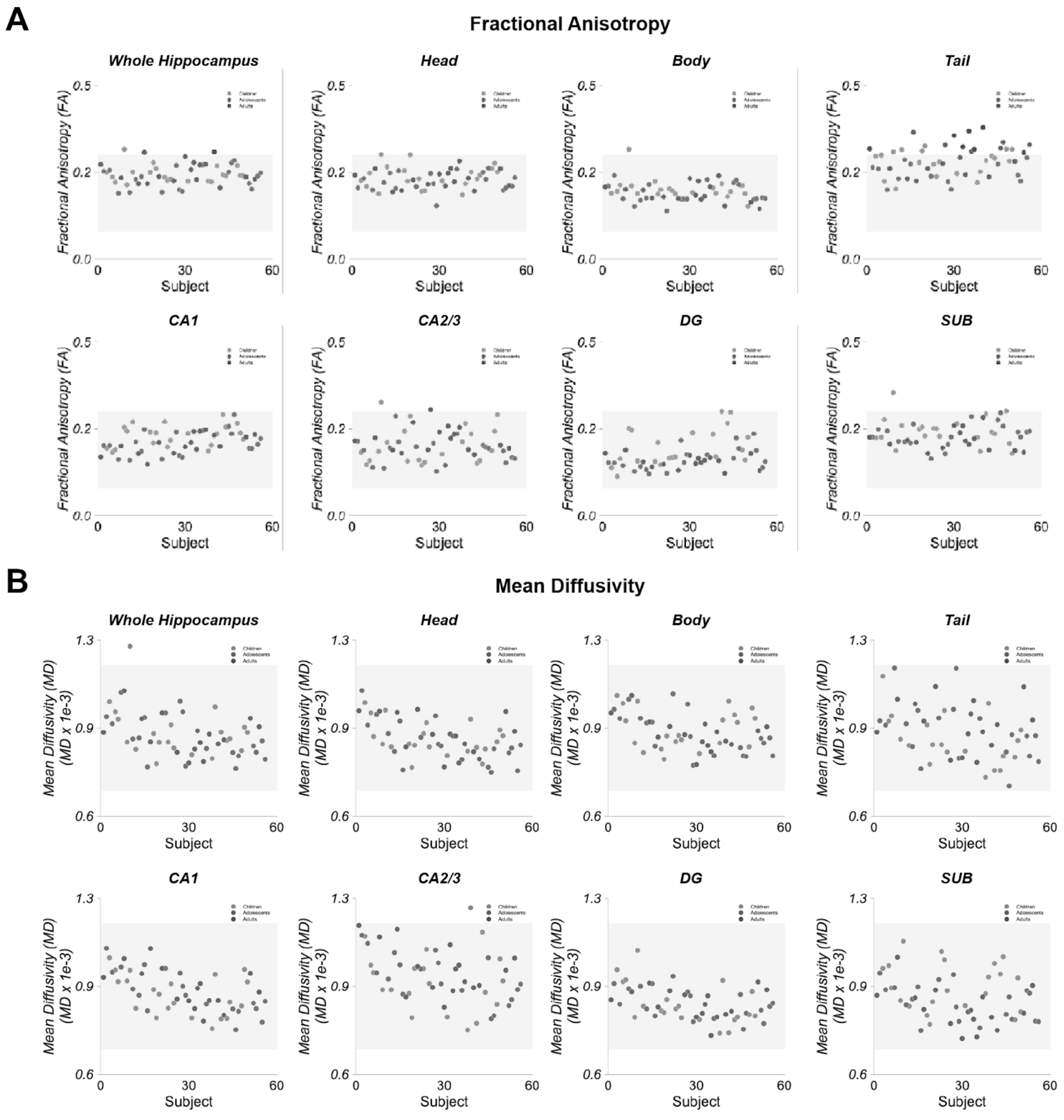


**Supplementary Figure 4.** Diffusion measurements for each subregion and subfield. **A.** Fractional anisotropy. **B.** Mean diffusivity. Gray areas represent the expected ranges.

***Supplementary analysis: Relationships between FA and data quality.***

We evaluated the relationship between fractional anisotropy (FA) and measures of data quality, framewise displacement (FD) (**Supplementary Figure 5A**) and signal-to-noise-ratio (SNR) (**Supplementary Figure 5B**). There was a relationship between SNR and FA in the CA2/3 subfield, $\beta = 0.0026$, $p = 0.03$. No other relationships were significant, all $p$s > 0.05.

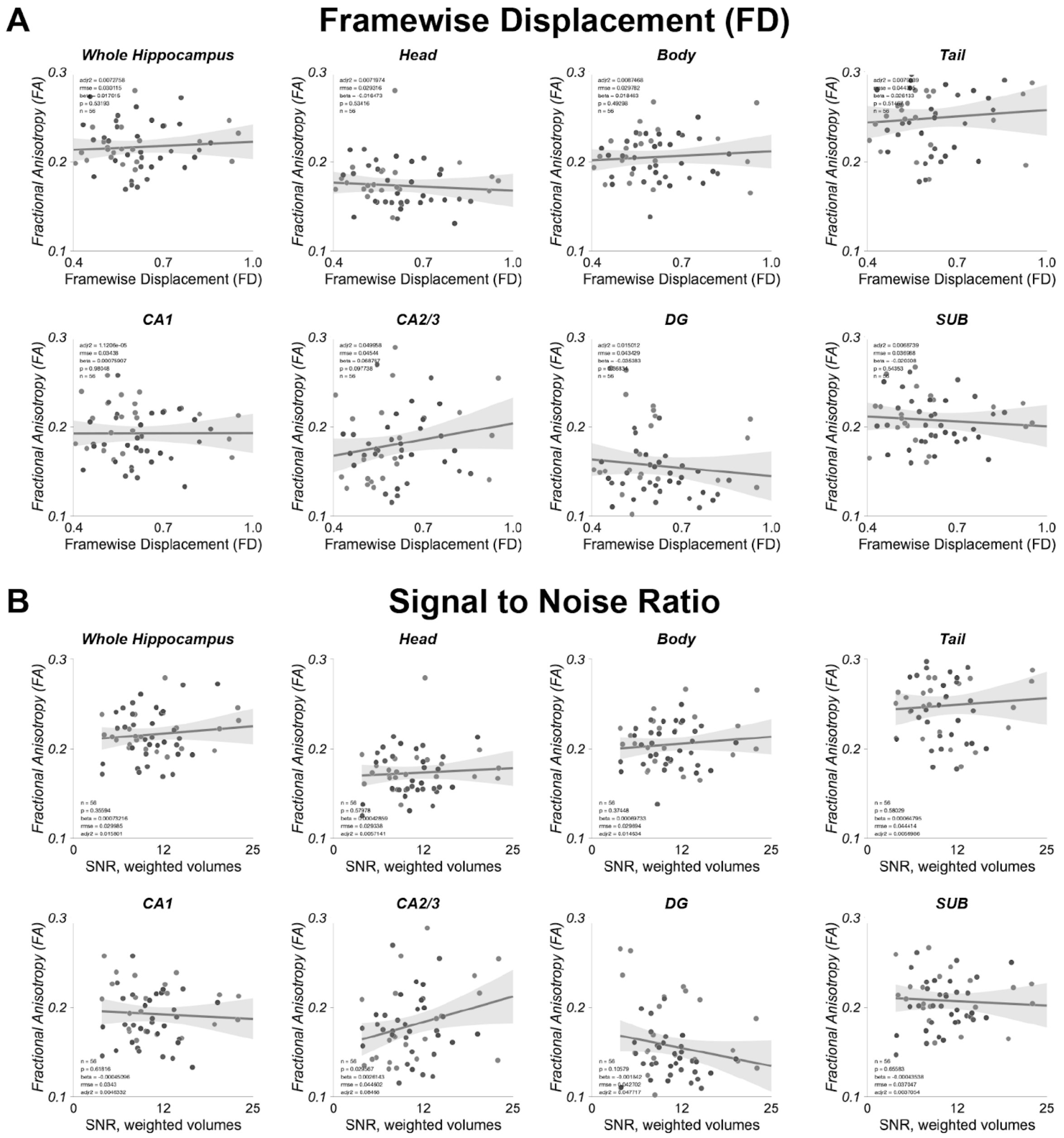


**Supplementary Figure 5.** Fractional Anisotropy. **A.** Framewise displacement. **B.** SNR, weighted volumes.

***Supplementary analysis: Relationships between MD and data quality.***

We evaluated the relationship between mean diffusivity (MD) and measures of data quality, framewise displacement (FD) (**Supplementary Figure 6A**) and signal-to-noise-ratio (SNR) (**Supplementary Figure 6B**). There was a relationship between FD and MD in the head subregion, $\beta$ = -0.171, $p$ = 0.011, in the CA2/3 subfield, $\beta$ = -0.220, $p$ = 0.032, and in the subiculum (SUB) subfield, $\beta$ = -0.216, $p$ = 0.008. No other relationships were significant, all $p$s > 0.05.

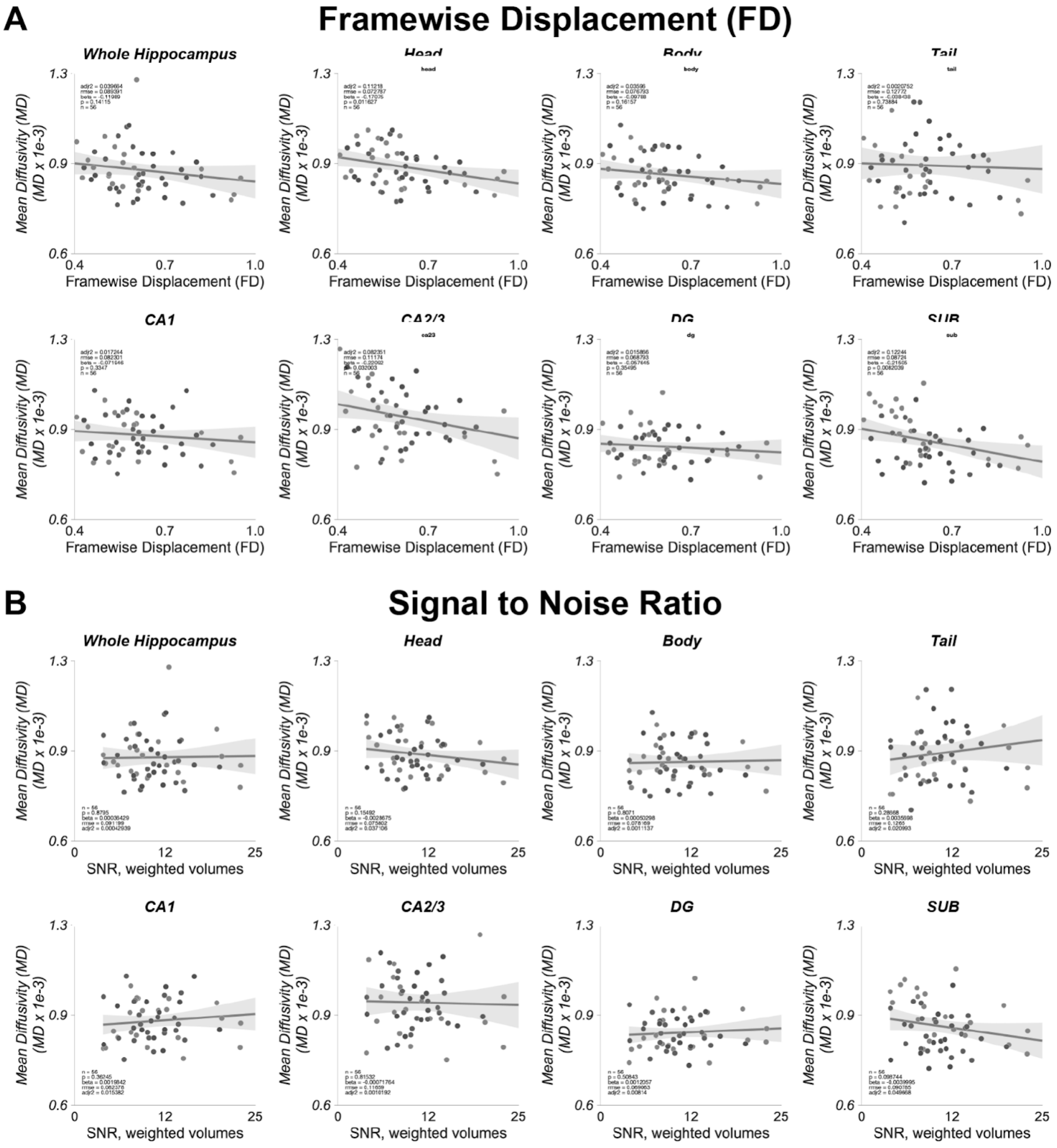


**Supplementary Figure 6.** Mean diffusivity. **A.** Framewise displacement. **B.** SNR, weighted volumes.

### Supplementary tables

### Supplementary Table 1.

#### Pearson correlations among variables of interest

| | | 1 | 2 | 3 | 4 | 5 | 6 | 7 | 8 | 9 | 10 | 11 | 12 |
|---|---|---|---|---|---|---|---|---|---|---|---|---|---|
| **1** | Age | 1.00 | | | | | | | | | | | |
| **2** | Head | -0.07 | 1.00 | | | | | | | | | | |
| **3** | Body | 0.04 | 0.14 | 1.00 | | | | | | | | | |
| **4** | Tail | 0.12 | **0.36** | **0.54*** | 1.00 | | | | | | | | |
| **5** | CA1 | -0.24 | **0.39** | **0.53*** | **0.60*** | 1.00 | | | | | | | |
| **6** | CA2/3 | 0.14 | 0.09 | **0.52*** | **0.33** | 0.15 | 1.00 | | | | | | |
| **7** | DG | **-0.42** | **0.27** | 0.03 | 0.05 | **0.38** | -0.03 | 1.00 | | | | | |
| **8** | SUB | -0.24 | **0.82*** | 0.06 | **0.32** | **0.44** | -0.24 | **0.40** | 1.00 | | | | |
| **9** | M Matching | **0.38** | -0.16 | -0.01 | 0.14 | -0.13 | 0.03 | -0.06 | -0.14 | 1.00 | | | |
| **10** | M Inference | **0.40** | -0.14 | 0.10 | 0.22 | -0.07 | 0.09 | **-0.31** | -0.20 | **0.74*** | 1.00 | | |
| **11** | P Matching | **0.45** | 0.02 | 0.08 | 0.27 | -0.10 | 0.10 | -0.16 | -0.01 | 0.27 | **0.30** | 1.0 | |
| **12** | P Inference | 0.20 | 0.06 | 0.07 | 0.11 | -0.12 | -0.03 | -0.18 | 0.04 | **0.31** | **0.33** | 0.19 | 1.0 |

*NOTE: Bolded values are significant at **p < 0.05**. Items with an asterisk remain significant after a Bonferonni correction for multiple comparisons, i.e., *** p = 0.05/66 correlations = 0.0007**. CA1 = cornu ammonis field 1; CA2/3 = cornu ammonis fields 2 and 3; DG = dentate gyrus; SUB = subiculum.*

**Supplementary Table 2.**

**Models tested for Hippocampal Subregion and Age Relationship**
**Fractional Anisotropy (FA)**

| *Subregion* | *Regression Model* | *N* | *AICc* | *Adj. $R^2$* | *F* | *p* |
|---|---|---|---|---|---|---|
| *Hippocampus* | hip ~ age | 52 | 231.94 | -0.016 | 0.190 | 0.665 |
| | hip ~ $age^2$ | 52 | 229.75 | -0.036 | 0.121 | 0.886 |
| | **hip ~ sex** | **52** | **234.98** | **0.049** | **3.62** | **0.063** |
| | hip ~ sex + age | 52 | 233.15 | 0.037 | 1.99 | 0.147 |
| | hip ~ sex + $age^2$ | 52 | 230.81 | 0.018 | 1.31 | 0.283 |
| | hip ~ sex * age | 54 | 232.75 | 0.034 | 1.61 | 0.198 |
| | hip ~ sex * $age^2$ | 54 | 227.97 | -0.003 | 0.967 | 0.447 |
| *Head* | head ~ age | 50 | 246.46 | 0.021 | 2.05 | 0.160 |
| | head ~ $age^2$ | 51 | 246.23 | 0.018 | 1.46 | 0.243 |
| | **head ~ sex** | **52** | **255.03** | **0.156** | **10.4** | **0.002** |
| | head ~ sex + age | 51 | 252.20 | 0.131 | 4.77 | 0.013 |
| | head ~ sex + $age^2$ | 53 | 252.37 | 0.106 | 3.05 | 0.037 |
| | head ~ sex * age | 51 | 252.66 | 0.160 | 4.18 | 0.011 |
| | head ~ sex * $age^2$ | 52 | 250.54 | 0.169 | 3.07 | 0.018 |
| *Body* | body ~ age | 52 | 236.56 | -0.019 | 0.048 | 0.828 |
| | body ~ $age^2$ | 51 | 234.66 | 0.009 | 1.22 | 0.305 |
| | **body ~ sex** | **53** | **237.64** | **-0.014** | **0.257** | **0.614** |
| | body ~ sex + age | 53 | 235.47 | -0.033 | 0.163 | 0.850 |
| | body ~ sex + $age^2$ | 52 | 233.28 | -0.010 | 0.836 | 0.481 |
| | body ~ sex * age | 51 | 232.19 | -0.015 | 0.758 | 0.523 |
| | body ~ sex * $age^2$ | 50 | 228.04 | 0.029 | 1.29 | 0.285 |
| *Tail* | **tail ~ age** | **55** | **188.35** | **-0.016** | **0.160** | **0.691** |
| | tail ~ $age^2$ | 54 | 185.93 | -0.025 | 0.351 | 0.706 |
| | tail ~ sex | 51 | 188.00 | -0.013 | 0.365 | 0.549 |
| | tail ~ sex + age | 54 | 187.07 | -0.025 | 0.363 | 0.697 |
| | tail ~ sex + $age^2$ | 54 | 185.14 | -0.037 | 0.366 | 0.778 |
| | tail ~ sex * age | 54 | 187.23 | 0.002 | 1.04 | 0.384 |
| | tail ~ sex * $age^2$ | 55 | 180.32 | -0.070 | 0.298 | 0.912 |

*Best fit models are bolded, selected based on AICc.*

**Supplementary Table 3.**

**Models tested for Hippocampal Subregion and Age Relationship**
**Mean Diffusivity (MD)**

| *Subregion* | *Regression Model* | *N* | *AICc* | *Adj. $R^2$* | *F* | *p* |
|---|---|---|---|---|---|---|
| *Hippocampus* | hip ~ age | 53 | 127.43 | -0.020 | 0.006 | 0.940 |
| | hip ~ $age^2$ | 53 | 125.42 | -0.035 | 0.114 | 0.892 |
| | **hip ~ sex** | **53** | **128.21** | **-0.005** | **0.757** | **0.388** |
| | hip ~ sex + age | 53 | 125.96 | -0.025 | 0.371 | 0.692 |
| | hip ~ sex + $age^2$ | 53 | 123.78 | -0.043 | 0.293 | 0.830 |
| | hip ~ sex * age | 53 | 123.68 | -0.044 | 0.263 | 0.851 |
| | hip ~ sex * $age^2$ | 52 | 119.12 | -0.059 | 0.461 | 0.803 |
| *Head* | head ~ age | 54 | 128.53 | 0.008 | 1.440 | 0.235 |
| | head ~ $age^2$ | 54 | 126.65 | -0.004 | 0.883 | 0.420 |
| | **head ~ sex** | **54** | **128.31** | **0.004** | **1.230** | **0.273** |
| | head ~ sex + age | 54 | 127.29 | 0.007 | 1.200 | 0.310 |
| | head ~ sex + $age^2$ | 54 | 125.19 | -0.008 | 0.860 | 0.468 |
| | head ~ sex * age | 55 | 123.64 | -0.024 | 0.580 | 0.631 |
| | head ~ sex * $age^2$ | 55 | 125.22 | 0.084 | 1.990 | 0.097 |
| *Body* | body ~ age | 55 | 127.91 | -0.014 | 0.250 | 0.619 |
| | body ~ $age^2$ | 55 | 126.06 | -0.026 | 0.307 | 0.737 |
| | body ~ sex | 55 | 127.93 | -0.014 | 0.272 | 0.604 |
| | body ~ sex + age | 55 | 125.89 | -0.030 | 0.226 | 0.798 |
| | body ~ sex + $age^2$ | 55 | 124.05 | -0.040 | 0.302 | 0.824 |
| | body ~ sex * age | 55 | 123.56 | -0.050 | 0.149 | 0.930 |
| | **body ~ sex * $age^2$** | **53** | **131.21** | **0.138** | **2.670** | **0.033** |
| *Tail* | **tail ~ age** | **51** | **105.41** | **0.134** | **8.74** | **0.005** |
| | tail ~ $age^2$ | 51 | 103.24 | 0.118 | 4.330 | 0.019 |
| | tail ~ sex | 51 | 97.06 | -0.020 | 0.017 | 0.897 |
| | tail ~ sex + age | 51 | 103.19 | 0.117 | 4.300 | 0.019 |
| | tail ~ sex + $age^2$ | 51 | 100.90 | 0.010 | 2.830 | 0.048 |
| | tail ~ sex * age | 51 | 101.17 | 0.104 | 2.930 | 0.043 |
| | tail ~ sex * $age^2$ | 51 | 96.477 | 0.070 | 1.760 | 0.141 |

*Best fit models are bolded, selected based on AICc.*

**Supplementary Table 4.**

**Models tested for Hippocampal Subfield and Age Relationship**
**Mean Diffusivity (MD)**

| *Subfield* | *Regression Model* | *N* | *AICc* | *Adj. $R^2$* | *F* | *p* |
|---|---|---|---|---|---|---|
| *CA1* | **CA1 ~ age** | **53** | **122.11** | **-0.010** | **0.485** | **0.490** |
| | CA1 ~ $age^2$ | 53 | 120.11 | -0.025 | 0.356 | 0.702 |
| | CA1 ~ sex | 53 | 121.86 | -0.007 | 0.659 | 0.421 |
| | CA1 ~ sex + age | 54 | 119.98 | -0.019 | 0.500 | 0.610 |
| | CA1 ~ sex + $age^2$ | 53 | 117.92 | -0.043 | 0.283 | 0.837 |
| | CA1 ~ sex * age | 52 | 119.14 | -0.027 | 0.553 | 0.649 |
| | CA1 ~ sex * $age^2$ | 51 | 116.33 | 0.007 | 1.070 | 0.388 |
| *CA23* | **CA23 ~ age** | **54** | **91.402** | **0.019** | **2.000** | **0.163** |
| | CA23 ~ $age^2$ | 54 | 90.204 | 0.019 | 1.500 | 0.233 |
| | CA23 ~ sex | 54 | 89.377 | -0.019 | 0.017 | 0.898 |
| | CA23 ~ sex + age | 54 | 89.157 | -0.001 | 0.983 | 0.381 |
| | CA23 ~ sex + $age^2$ | 54 | 87.884 | -0.001 | 0.987 | 0.407 |
| | CA23 ~ sex * age | 53 | 88.590 | -0.021 | 0.641 | 0.593 |
| | CA23 ~ sex * $age^2$ | 53 | 84.810 | -0.041 | 0.595 | 0.704 |
| *DG* | **DG ~ age** | **53** | **148.00** | **0.007** | **1.360** | **0.248** |
| | DG ~ $age^2$ | 53 | 146.47 | 0.001 | 1.020 | 0.369 |
| | DG ~ sex | 53 | 147.06 | -0.011 | 0.437 | 0.512 |
| | DG ~ sex + age | 53 | 146.09 | -0.007 | 0.830 | 0.442 |
| | DG ~ sex + $age^2$ | 53 | 144.35 | -0.016 | 0.736 | 0.536 |
| | DG ~ sex * age | 53 | 143.78 | -0.027 | 0.552 | 0.649 |
| | DG ~ sex * $age^2$ | 52 | 142.69 | 0.033 | 1.350 | 0.261 |
| *SUB* | **SUB ~ age** | **53** | **121.91** | **0.170** | **11.600** | **0.001** |
| | SUB ~ $age^2$ | 53 | 120.53 | 0.167 | 6.220 | 0.004 |
| | SUB ~ sex | 51 | 115.61 | 0.030 | 2.560 | 0.116 |
| | SUB ~ sex + age | 53 | 121.33 | 0.180 | 6.690 | 0.003 |
| | SUB ~ sex + $age^2$ | 52 | 120.74 | 0.228 | 6.010 | 0.002 |
| | SUB ~ sex * age | 53 | 118.99 | 0.163 | 4.370 | 0.008 |
| | SUB ~ sex * $age^2$ | 52 | 120.45 | 0.333 | 6.090 | 0.002 |

*Best fit models are bolded, selected based on AICc.*

**Supplementary Table 5.**

**Models tested for Hippocampal Subfield and Age Relationship**
**Subregion: Head**
**Fractional Anisotropy (FA)**

| *Subregion* | *Subfield* | *Regression Model* | *N* | *AICc* | *Adj. $R^2$* | *F* | *p* |
|---|---|---|---|---|---|---|---|
| *Head* | *CA1* | CA1 ~ age | 52 | 231.93 | 0.146 | 9.710 | 0.003 |
| | | CA1 ~ $age^2$ | 50 | 230.99 | 0.302 | 11.600 | 0.0001 |
| | | CA1 ~ sex | 53 | 230.32 | 0.114 | 7.720 | 0.008 |
| | | CA1 ~ sex + age | 52 | 235.80 | 0.286 | 11.200 | 0.0001 |
| | | CA1 ~ sex + $age^2$ | 52 | 233.85 | 0.286 | 7.820 | 0.0002 |
| | | CA1 ~ sex * age | 52 | 233.90 | 0.278 | 7.530 | 0.0003 |
| | | **CA1 ~ sex * $age^2$** | **49** | **245.19** | **0.544** | **12.500** | **0.0000002** |
| *Head* | *CA23* | CA2/3 ~ age | 52 | 226.78 | -0.019 | 0.047 | 0.829 |
| | | CA2/3 ~ $age^2$ | 54 | 226.59 | 0.025 | 1.690 | 0.196 |
| | | CA2/3 ~ sex | 53 | 227.393 | -0.015 | 0.211 | 0.648 |
| | | CA2/3 ~ sex + age | 54 | 225.322 | -0.028 | 0.279 | 0.757 |
| | | CA2/3 ~ sex + $age^2$ | 55 | 225.542 | 0.072 | 2.400 | 0.079 |
| | | CA2/3 ~ sex * age | 56 | 227.160 | 0.034 | 1.650 | 0.189 |
| | | **CA2/3 ~ sex * $age^2$** | **52** | **227.399** | **0.282** | **5.000** | **0.001** |
| *Head* | *DG* | DG ~ age | 48 | 233.04 | 0.026 | 2.270 | 0.139 |
| | | DG ~ $age^2$ | 48 | 234.061 | 0.071 | 2.790 | 0.072 |
| | | DG ~ sex | 48 | 237.567 | 0.047 | 3.330 | 0.075 |
| | | DG ~ sex + age | 47 | 237.341 | 0.094 | 3.390 | 0.043 |
| | | **DG ~ sex + $age^2$** | **48** | **236.586** | **0.142** | **3.590** | **0.021** |
| | | DG ~ sex * age | 47 | 235.931 | 0.092 | 2.560 | 0.067 |
| | | DG ~ sex * $age^2$ | 48 | 226.085 | 0.185 | 3.130 | 0.017 |
| *Head* | *SUB* | SUB ~ age | 51 | 229.277 | 0.015 | 1.780 | 0.188 |
| | | SUB ~ $age^2$ | 51 | 227.019 | -0.005 | 0.873 | 0.424 |
| | | **SUB ~ sex** | **53** | **229.365** | **0.016** | **1.840** | **0.180** |
| | | SUB ~ sex + age | 53 | 228.806 | 0.028 | 1.740 | 0.185 |
| | | SUB ~ sex + $age^2$ | 53 | 226.810 | 0.014 | 1.250 | 0.301 |
| | | SUB ~ sex * age | 52 | 228.542 | 0.096 | 2.800 | 0.050 |
| | | SUB ~ sex * $age^2$ | 51 | 226.945 | 0.237 | 2.800 | 0.023 |

*Best fit models are bolded, selected based on AICc.*

**Supplementary Table 6.**

**Models tested for Hippocampal Subfield and Age Relationship**
**Subregion: Body**
**Fractional Anisotropy (FA)**

| *Subregion* | *Subfield* | *Regression Model* | *N* | *AICc* | *Adj. $R^2$* | *F* | *p* |
|---|---|---|---|---|---|---|---|
| *Body* | *CA1* | CA1 ~ age | 53 | 227.023 | 0.204 | 14.400 | 0.0004 |
| | | **CA1 ~ $age^2$** | **50** | **232.243** | **0.426** | **19.2** | **0.000001** |
| | | CA1 ~ sex | 53 | 217.951 | 0.203 | -0.016 | 0.654 |
| | | CA1 ~ sex + age | 53 | 224.790 | 0.189 | 7.050 | 0.002 |
| | | CA1 ~ sex + $age^2$ | 50 | 230.029 | 0.415 | 12.600 | 0.000004 |
| | | CA1 ~ sex * age | 53 | 228.176 | 0.296 | 8.280 | 0.0002 |
| | | CA1 ~ sex * $age^2$ | 53 | 232.740 | 0.387 | 7.560 | 0.00003 |
| *Body* | *CA23* | CA2/3 ~ age | 49 | 164.558 | 0.348 | 26.600 | 0.000005 |
| | | CA2/3 ~ $age^2$ | 49 | 162.296 | 0.334 | 13.000 | 0.00003 |
| | | CA2/3 ~ sex | 53 | 148.479 | 0.005 | 1.250 | 0.270 |
| | | **CA2/3 ~ sex + age** | **50** | **165.134** | **0.346** | **14.000** | **0.00002** |
| | | CA2/3 ~ sex + $age^2$ | 50 | 162.845 | 0.333 | 9.160 | 0.00007 |
| | | CA2/3 ~ sex * age | 50 | 162.823 | 0.333 | 9.150 | 0.00007 |
| | | CA2/3 ~ sex * $age^2$ | 51 | 156.536 | 0.261 | 4.530 | 0.002* |
| *Body* | *DG* | **DG ~ age** | **49** | **213.579** | **0.016** | **1.760** | **0.191** |
| | | DG ~ $age^2$ | 49 | 212.642 | 0.021 | 1.520 | 0.229 |
| | | DG ~ sex | 50 | 212.684 | -0.013 | 0.360 | 0.551 |
| | | DG ~ sex + age | 49 | 211.336 | -0.005 | 0.876 | 0.423 |
| | | DG ~ sex + $age^2$ | 49 | 210.266 | -0.001 | 0.992 | 0.405 |
| | | DG ~ sex * age | 49 | 209.604 | -0.014 | 0.777 | 0.513 |
| | | DG ~ sex * $age^2$ | 49 | 206.740 | -0.014 | 0.866 | 0.512 |
| *Body* | *SUB* | SUB ~ age | 53 | 180.980 | -0.016 | 0.159 | 0.692 |
| | | SUB ~ $age^2$ | 52 | 183.500 | 0.064 | 2.730 | 0.075 |
| | | SUB ~ sex | 53 | 181.099 | -0.014 | 0.274 | 0.603 |
| | | SUB ~ sex + age | 53 | 178.974 | -0.032 | 0.194 | 0.825 |
| | | **SUB ~ sex + $age^2$** | **52** | **181.642** | **0.053** | **1.950** | **0.134** |
| | | SUB ~ sex * age | 53 | 177.189 | -0.042 | 0.301 | 0.825 |
| | | SUB ~ sex * $age^2$ | 52 | 177.767 | 0.033 | 1.350 | 0.260 |

*Best fit models are bolded, selected based on AICc.*
